# Sniffing the Human Body-Volatile Hexadecanal Blocks Aggression in Men but Triggers Aggression in Women

**DOI:** 10.1101/2020.09.29.318287

**Authors:** Eva Mishor, Daniel Amir, Tali Weiss, Danielle Honigstein, Aharon Weissbrod, Ethan Livne, Lior Gorodisky, Shiri Karagach, Aharon Ravia, Kobi Snitz, Diyala Karawani, Rotem Zirler, Reut Weissgross, Timna Soroka, Yaara Endevelt-Shapira, Shani Agron, Liron Rozenkrantz, Netta Reshef, Edna Furman-Haran, Heinz Breer, Joerg Strotmann, Noam Sobel

## Abstract

Body-volatiles can effectively trigger or block conspecific aggression in terrestrial mammals. Here we tested whether hexadecanal (HEX), a human body-volatile implicated as a mammalian-wide social cue, impacts human aggression. Using validated behavioural paradigms, we observed a remarkable dissociation: sniffing HEX blocked aggression in men, but triggered aggression in women. Next, using functional brain imaging, we uncovered a pattern of brain activity mirroring behaviour: In both men and women, HEX increased activity in the left angular gyrus, an area implicated in perception of social cues. Hex then modulated functional connectivity between the angular gyrus and a brain network implicated in social appraisal (temporal pole) and aggressive execution (amygdala and orbitofrontal cortex) in a sex-dependent manner consistent with behaviour: increasing connectivity in men, but decreasing connectivity in women. These findings implicate sex-specific social chemosignaling at the mechanistic heart of human aggressive behaviour.

## Introduction

Terrestrial mammalian aggressive behavior can be triggered or blocked by social odors (*1–4*). For example, a rabbit mother will attack and even kill her pups if they are tainted with the body-odor of a stranger female (*5*), and two specific volatile components in mouse urine trigger fighting between males (*6*). Such aggression-triggering chemosignals are not all volatile: specific major urinary proteins (MUPs) can also trigger aggression, an effect mediated by the accessory olfactory system (*7*). In turn, chemosignals not only trigger aggression, they can also block it. For example, tainting a pig with the known volatile reproductive pheromone androstenone (5a-Androst-16-en-3-one) blocks conspecific aggression towards the tainted pig (*8*). Also, a mouse tear-bound peptide blocks male mouse aggression towards mouse pups tainted with the peptide (*9*). Such tear-bound aggression-blocking chemosignals may be a mammalian-wide phenomenon: male mole-rats cover themselves with their own harderian secretions, and this blocks aggression towards them from dominant male conspecifics (*10*). Relatedly, sniffing human tears reduces testosterone in men (*11, 12*), but the impact of this on aggressive behavior has yet to be investigated. To conclude, both volatile and non-volatile chemosignals can either trigger or block mammalian aggression.

Like all mammals, humans engage in reactive aggression from a very early age (*13*), and throughout life (*14*). Although aggression is a major factor in the human condition, the mechanistic neural substrates of human aggression are not well-understood (*15–17*). Could human aggression mechanisms be tied to chemosignaling mechanisms as they are in all other terrestrial mammals? Two studies have suggested that humans emit aggression-specific body-odors (*18, 19*), but whether and how human aggressive behaviour is then impacted by social chemosignals remains unknown. To test the hypothesis that a social chemosignal can modulate human aggression through modulation of neural activity we used standard behavioural and neuroimaging aggression paradigms with and without a concurrent social chemosignal. As a social chemosignal, we tested Hexadecanal (HEX), a volatile long-chain aliphatic aldehyde, first identified as a social-buffering agent in mice (*20*). Although species-specificity has been considered a hallmark of social chemosignaling (*21*), some signaling molecules may be conserved across species, with either similar or different signal meaning (*22, 23*). Because the mouse receptor for HEX (OR37B) is highly conserved across mammals, it has been suggested that HEX may be one such cross-species conserved signaling molecule (*24*). Humans indeed express an OR37 receptor orthologue (*25, 26*), and indeed emit HEX in feces, skin, and breath (*27, 28*). Moreover, exposure to HEX reduced startle-responses in humans (*29*), implicating impact on arousal. With all this in mind, we tested whether and how HEX impacts human aggression.

## Results

### HEX blocked aggression in men but triggered aggression in women

To gauge aggression, we used a modified Taylor aggression paradigm (TAP), a well-established measure of human aggressive behaviour (*30–33*). A total of 127 participants (67 men, mean age 25.48 ± 3.46, range 21-34) played the game in a double-blind between-subjects design, half concurrent with exposure to HEX obscured in carrier (Eugenol), and half concurrent with exposure to carrier alone (Figure 1A). To ask whether the addition of HEX was associated with an explicit percept, participants used a visual-analogue scale (VAS) to rate the stimuli along the primary dimensions of human olfactory perception; namely intensity and pleasantness (*34*) (Figure 1B, 1C). Because odorant ratings did not distribute normally (Pleasantness: Shapiro-Wilk W = 0.98, p = 0.001, Intensity: W = 0.96, p = 0.002) we applied a linear mixed model with factors of Sex and Odor and random effects of participant, which revealed that HEX had no impact on perceived odorant pleasantness (Sex: F(1, 123) = 1.13, p = 0.29, Odor: F(1, 123) = 0.12, p = 0.73, Sex x Odor: F(1, 123) = 0.10, p = 0.7), and no overall impact on perceived intensity, but a marginal yet statistically significant interaction (Sex: F(1, 123) = 3.66, p = 0.06, Odor: F(1, 123) = 0.6, p = 0.44, Sex x Odor: F(1, 123) = 6.07, p = 0.02). Post-hoc comparisons (Tukey adjustment for multiple comparisons), however, revealed that this interaction reflected opposing insignificant effects in men and women (Men Control: mean VAS = −0.09 ± 0.07, men HEX: mean VAS = −0.22 ± 0.77, t(123) = 1.10, p = 0.69. Women Control: mean VAS = −0.09 ± 0.60, women HEX: mean VAS = 0.38 ± 0.96, t(123) = 2.35, p = 0.09) (Figure 1D, 1E). In sum, the addition of HEX to the carrier had no impact on perceived stimulus pleasantness, and minimal impact on perceived stimulus intensity.

**Figure 1.**
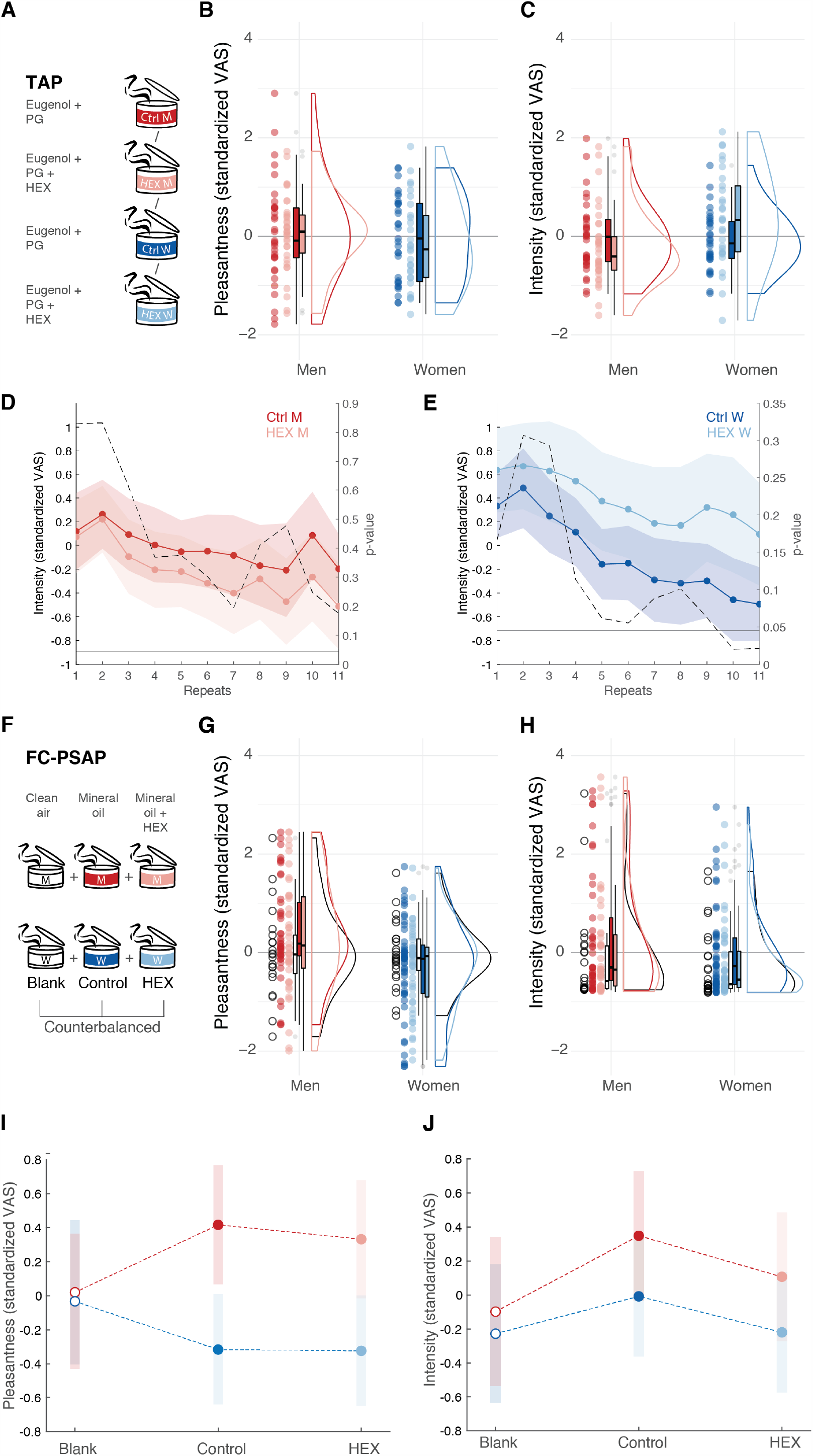
HEX did not significantly shift stimulus perception. (**A**) The between-subjects TAP included four groups exposed to either control (100 ul, 10% eugenol in propylene glycol (PG)) or HEX (100 ul, 0.083M HEX in 10% eugenol in PG). Control n = 34M and 31F, HEX n = 33M and 29F. (**B**) TAP odorant pleasantness ratings along the VAS. Each dot is a participant, the thick horizontal line is the median, the rectangle reflects the interquartile range (25th to the 75th percentiles) and the whiskers are no more than 1.5 * IQR of the upper and lower hinges. Outlying points are plotted individually. (**C**) TAP odorant Intensity ratings along the VAS. Elements as in B. (**D**) Mean Intensity and confidence interval for 11 consecutive exposures to HEX and Control, in men. The dotted line is the point-by-point p-value for the HEX-Control t-test. The horizontal line represents the significance threshold p-value, corrected for multiple comparisons (Bonferroni correction), set to 0.0045. (**E**) Same as in D, in women. (**F**) The within-subjects FC-PCAP included two groups: men (n = 25) and women (n = 24). Both were exposed to Blank (clean air) Control (mineral oil) and HEX (0.083M HEX dissolved in mineral oil). (**G**) FC-PCAP odorant pleasantness ratings along the VAS. Elements as in B. (**H**) FC-PCAP odorant intensity ratings along the VAS. Elements as in B. (**I**) FC-PCAP odorant pleasantness ratings mean and confidence interval. (**J**) FC-PCAP odorant intensity ratings mean and confidence interval.

The TAP begins with a *provocation phase* where the participant plays an on-screen version of the *ultimatum game* (*35*) against a purported game-partner who is in fact a game algorithm (Figure 2A). In each round of the ultimatum game, both “players” are allotted a sum of money (∼$9) that they can keep if they agree on how to distribute it between them. The game is programmed such that the purported partner agrees only to distributions that significantly discriminate against the participant. Five such rounds served as an effective provocation. At this point, there were no differences between the groups (Supplementary Figure 1; Supplementary Figure 2). Following this, is an *aggression discharge phase* where the participant is again misled to believe they are playing against the same partner (who previously provoked them), but now in a reaction-time task where they compete to identify a change in shape of a target. The first to react is then allowed to blast his/her opponent with a loud noise blast. The volume applied by the participant is taken as a measure of aggression (*36, 37*) (Figure 2B). The game was rigged such that the participant was faster than the fictitious opponent on 16 of 27 rounds, and in rounds where the fictitious opponent was faster, the participant endured noise-blasts randomly ranging in volume.

**Figure 2.**
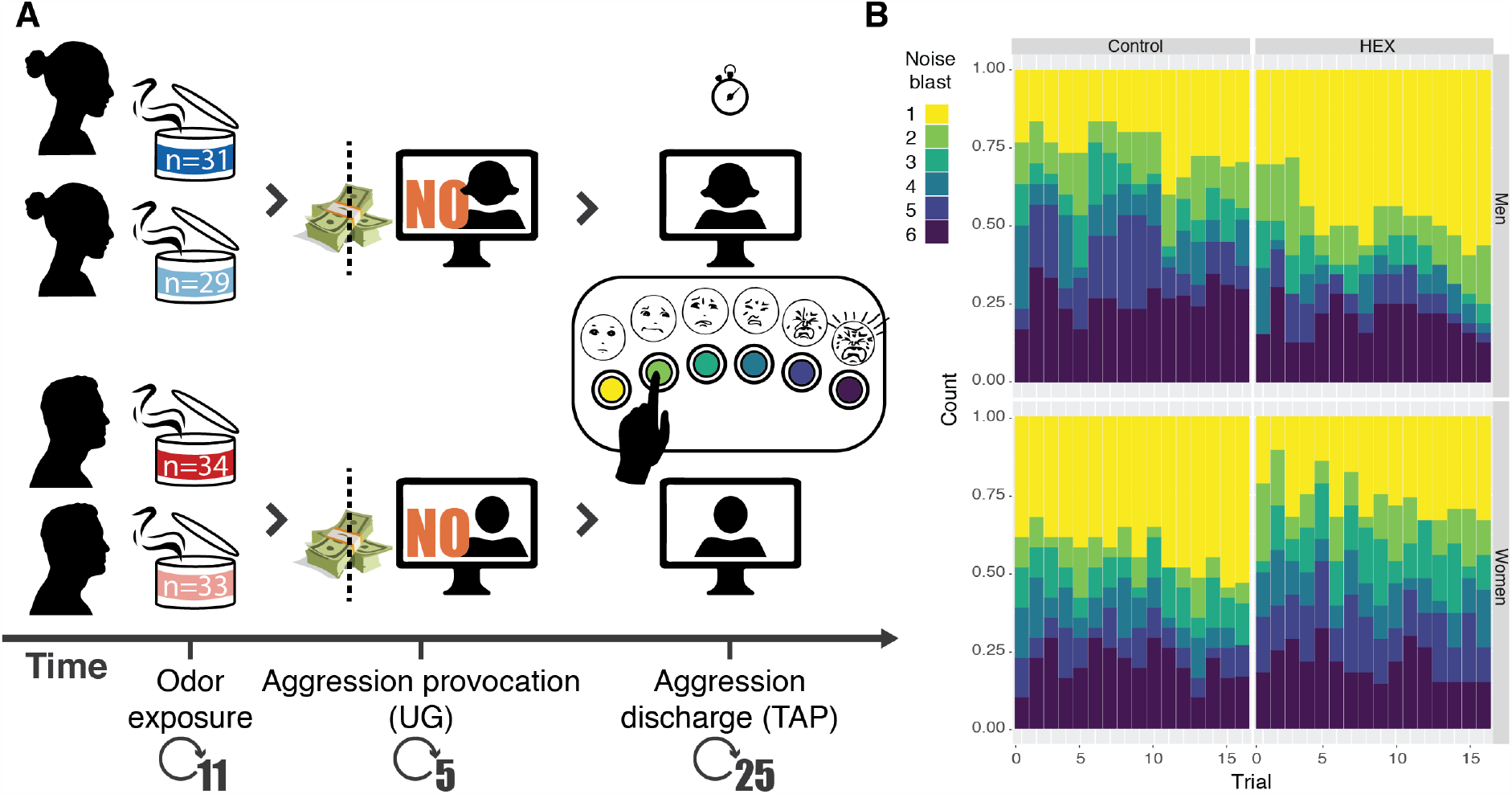
Path to provoking and gauging human aggressive behaviour. (**A**) In a between-subjects design, participants were exposed to an odorant (HEX or control), then played a game where their online partner was unfair towards them in monetary distribution (provocation), and then another game where they could blast that same (non-existent) person with noise-blasts (aggression discharge). (**B**) The complete distribution of noise-blasts applied in the study by men (n = 67) and women (n = 60) under HEX or control (yellow = mild, purple = harsh).

Noise-blast volume was entered into a repeated-measures ordinal logistic regression analysis with factors of Odorant (HEX/Control) and Sex (men/women), and random effect of Participant. We observed a trend towards an effect of Odorant (Z = 1.84, p = 0.07), no effect of Sex (Z = 1.63, p = 0.10), and a significant Odorant X Sex interaction (Z =2.03, p = 0.04, effect size: generalized η^2^ = 0.04). This significant interaction reflected that HEX significantly lowered aggression in men (men Hex: n = 34, mean blast vol = 2.76 ± 2.03, men Control: n = 33, mean blast vol = 3.60 ± 1.98, Mann-Whitney (trials compacted as repeated measure), Z = 5.855, p < 0.0001, effect size r = 0.7), yet significantly increased aggression in women (women Hex: n = 29, mean blast vol = 3.40 ± 1.91, women Control: n = 31, mean blast vol = 2.96±2.03, Mann-Whitney (trials compacted as repeated measure), Z = 3.655, p = 0.0003, effect size r = 0.47) (Figure 3A).

**Figure 3.**
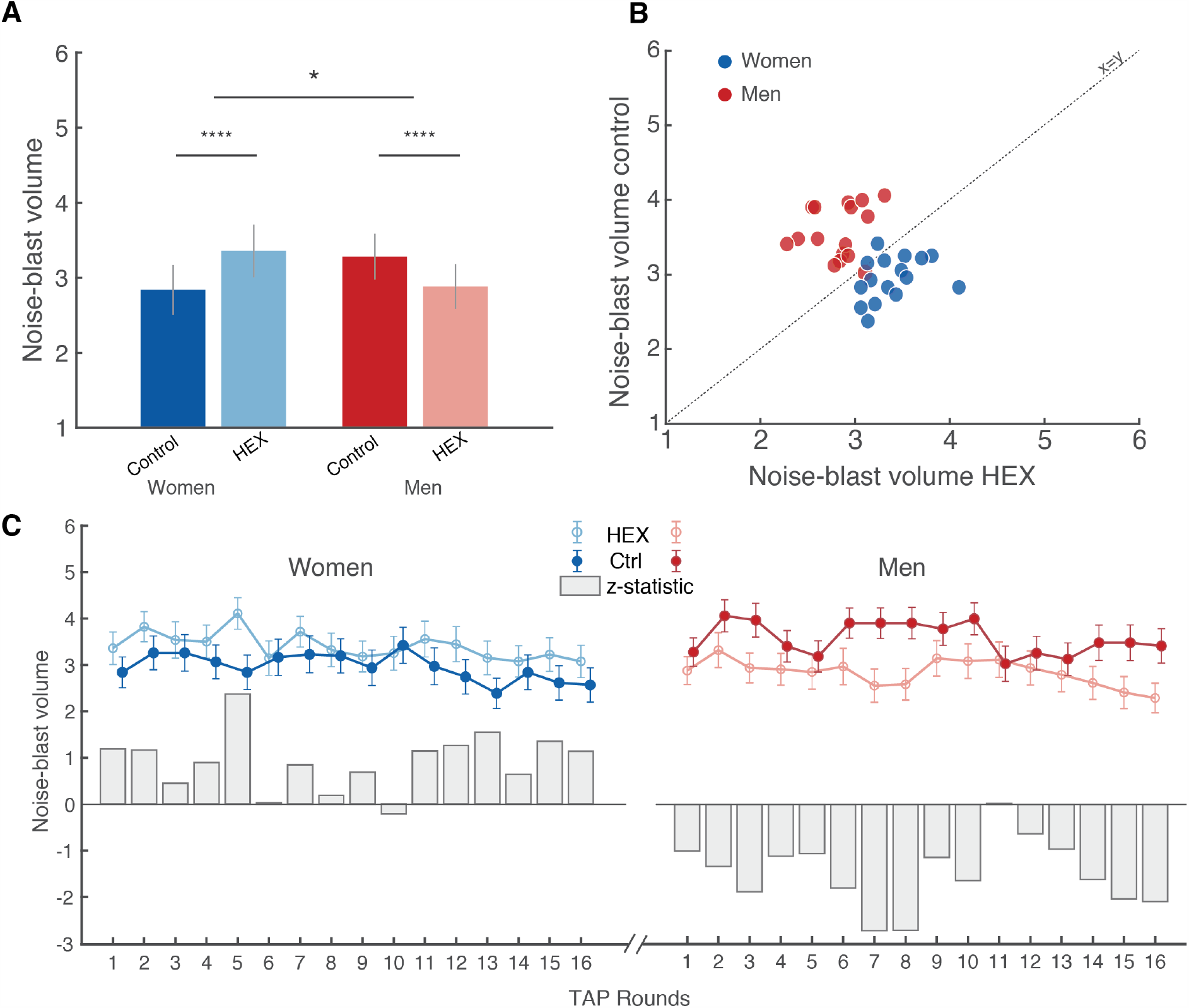
Exposure to HEX modulated aggressive behaviour in a sex-dependent manner across participants. (**A**) Group-mean noise-blast volume. (**B**) Trial-wise mean noise-blast volume. Each dot is the mean of a trial (1-16) of all men (red) or women (blue) participants. The Y axis reflects the trial mean during Control, and the X axis reflects the trial mean during HEX. The dotted line is the unit slope line (X=Y), such that data under the line reflect increased aggression in HEX and data above the line reflect decreased aggression in HEX. (**C**) A trial-by-trial depiction across participants. The solid dots reflect Control, the open circles reflect HEX and the grey bars reflect a trial-by-trial Z test. Men: n = 67 (34 Control). Women: n = 60 (31 Control). All tests were two-tailed, all error bars represent SEM. * = p < 0.05, **** = p < 0.0005.

To gain further insight into the dynamics of aggressive behaviour in the TAP, we looked at the 16 repeated trials separately. When depicting the trial-by-trial mean, there is an apparent small yet highly consistent difference between the conditions in both sexes (Figure 3B, 3C). To again quantify this, we compared the trial-by-trial means using a Mann-Whitney test, which provided a trial-wise Z value. An independent t-test on the trial-by-trial z-values again revealed a significant effect of condition within sex: significantly decreasing aggression in men exposed to HEX (Trial-wise blast volume: control = 3.57 ± 0.34, HEX = 2.91 ± 0.39, t(16) = −6.37, p<10^−5^) yet significantly increasing it in women (Trial-wise blast volume: control = 2.84 ± 0.55, HEX = 3.34 ± 0.33, t(15) = 4.70, p<0.0003) (these numbers differ from the previous analysis because they reflect trial-wise blast volume, whereas the previous analysis reflected participant-wise blast volume), again indicating a significant dissociation between the two sexes (t(15) = −11.18, p<10^−7^) (these differences again re-emerged in a validation permutation test (Supplementary Figure 3)). Behavioural differences were not associated with perceptual valence differences between the two conditions (Figure 1), were not related to vigilance (Supplementary Figure 4), and were not reflected in group-levels of Cortisol or Testosterone (Supplementary Figure 5). In sum, three separate analyses of the TAP implied that exposure to HEX reduced aggression in men but increased aggression in women.

We next used functional magnetic resonance imaging (fMRI) in order to ask where and how in the brain does HEX impact aggression. We used the point subtraction aggression paradigm (PSAP), another well-established paradigm in aggression research that has been used in fMRI (*38–40*). It allows imaging of brain activity during provocation (having your money taken away from you) and aggression (subtracting money from others, yet not for monetary profit). In order to provide for a more naturalistic aggressive output, we substituted the response buttons with fist-clench pressure sensors. Participants were misled to think that the response-device provides binary output alone, but in fact we measured fist-clench effort as a naturalistic implicit continuous measure of aggression extent. In support of this method, we observed a modest but significant correlation between measured aggression at baseline and basal aggressive tendencies as estimated by a standard aggression questionnaire (AGQ) (*41*) (r = 0.30, p = 0.04) (Supplementary Figure 6). Such correlations typically emerge only in very large samples (*42*), so their emergence here underlies the power of the fist-clench PSAP (FC-PSAP). In a within-subjects, double-blind design (Figure 1F), 49 participants (24 women, mean age 26.98 ± 3.92, range 19-36) completed an average of 134.39 (± 15.33) trials (of all types), about half concurrent with HEX and about half with carrier (mineral oil) alone, counter-balanced for order (For full table of events, see Supplementary Table 1).

**Figure 6.**
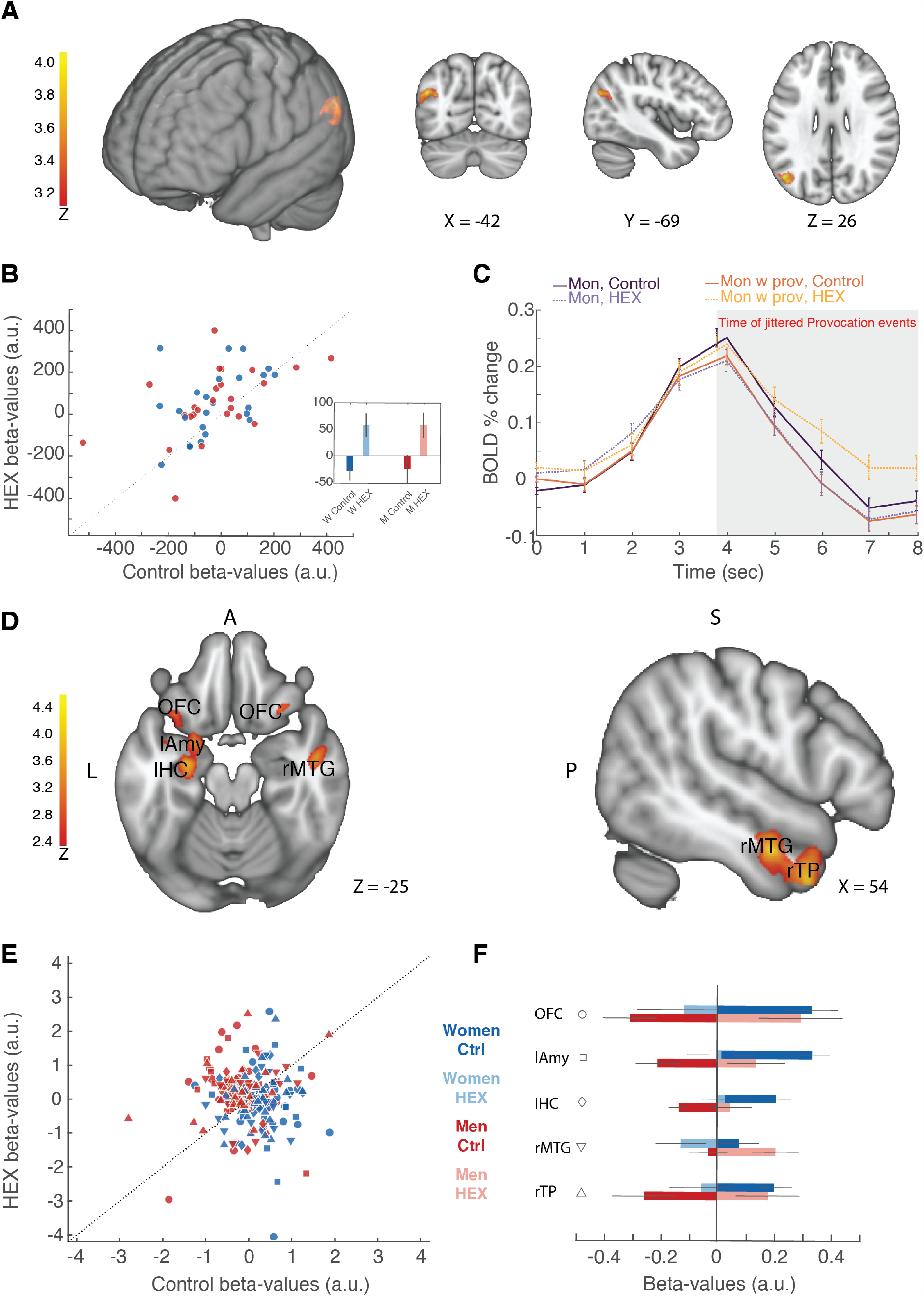
HEX drives sex-specific functional connectivity in the brain substrates of aggression. **(A)** A contrast ANOVA statistical parametric map depicting activity greater during provocation vs. baseline in HEX vs. Control (p < 0.001, corrected). **(B)** Beta-values extracted from the left angular gyrus. Each dot is a woman (blue) or man (red) participant. The Y axis reflects that participant’s mean Beta during HEX, and the X axis reflects that participant’s mean Beta during Control, during a Provocation. The dotted line is the unit slope line (X=Y), such that data under the line reflect increased activation in Control and data above the line reflect increased activation in HEX. (**C**) The percent signal change in BOLD activity in the left angular gyrus ROI, depicted by condition. (**D**) A contrast statistical parametric map depicting functional connectivity with the left angular gyrus greater during provocation vs. baseline in HEX vs. Control in Men > Women (p < 0.01, corrected). OFC – Orbitofrontal cortex, lAmy – left Amygdala, lHC – left Hippocampus, rMTG – right Medial temporal gyrus, rTP – right temporal pole. (**E**) Beta-values for functional connectivity with the angular gyrus. Each dot is a woman (blue) or man (red) participant, shape of dot is as depicted in panel F. (**F**) Bar graph of the beta values of functional connectivity of Provocation>Baseline, HEX>Control, Women>Men, for whole-brain connectivity analysis with the AG as a seed region.

### HEX again blocked aggression in men but triggered aggression in women

To again ask whether the addition of HEX was associated with an explicit percept, participants used a visual-analogue scale to rate the stimuli along the primary dimensions of human olfactory perception; namely intensity and pleasantness (*34*) (Figure 1G, 1H). Because odorant ratings again did not distribute normally (Pleasantness: W = 0.98, p = 0.001, intensity: W = 0.75, p = 2.2*10^−6^), we applied a linear mixed model with factors of Sex and Odor and random effects of participant. Analysis of the pleasantness ratings revealed a significant difference between the sexes in overall pleasantness in the FC-PSAP, no effect of Odor, and a marginally significant interaction (Sex: F(1, 43) = 8.07, p = 0.007, Odor: F(2, 175) = 0.007, p = 0.94, Sex x Odor: F(2, 175) = 3.08, p = 0.05). Post-hoc pairwise comparisons (Tukey adjustment for multiple comparisons) revealed that this difference arose from men’s ratings of the odorless control. In other words, this reflected a basic sex-difference in reported valence (men Control-women Control: men mean VAS = 0.3 ± 1.05, women mean VAS = −0.26 ± 0.87, t(43) = 3.09, p = 0.04. Men Control-women HEX: men mean VAS = 0.3 ± 1.05, women mean VAS = −0.32 ± 0.83, t(43) = 3.12, p = 0.04. All other comparisons non-significant (all t < 2.78, all p > 0.09) (Figure 1I). To conclude, there were no differences within sex, and a marginal difference across sexes arising from men’s ratings of the odorless control. The same model applied to intensity ratings reveled no main effect of sex, no interaction, but an effect of Odor (Sex: F(1, 43) = 1.69, p = 0.2, Odor: F(2, 175) = 3.88, p = 0.02, Sex x Odor: F(2, 175) = 0.41, p = 0.66). Post-hoc contrasts (Tukey adjustment for multiple comparisons) revealed that the intensity differences in the FC-PSAP arose from Control-Blank comparisons (blank mean VAS = −0.17 ± 0.88, control mean VAS = 0.15 ± 1.08, t(175) = 2.57, p = 0.03), but not Control-HEX comparisons (control mean VAS = 0.15 ± 1.08, HEX mean VAS = −0.07 ± −0.96, t(175) = 2.14, p = 0.08) (Figure 1J). In other words, mineral oil is detectable versus blank, but HEX had no impact on perception in this experiment.

In the PSAP, aggressive behaviour is typically reported as the aggression/provocation ratio (APR), namely the aggressive responses divided by the number of provocation events participants experienced. Higher APR values imply increased aggression. Using the within-subjects FC-PSAP, we observed the same behavioral pattern we previously observed using the between-subjects TAP (Figure 4C). More specifically, a Kruskal-Wallis test on the delta APR with a factor of Sex revealed a significant effect of Sex between odour conditions (χ2 = 9.24, df = 1, p = 0.002, effect size η^2^ = 0.18). This interaction reflected that HEX drove a significant reduction in aggression by men (APR control = 1.12 ± 1.43, APR HEX = 0.88 ± 1.08, Wilcoxon signed rank test: Z = 2.18, p = 0.03, effect size r = 0.45), yet increased aggression by women [we note that two women were 3 standard deviations away from the APR mean, one of them from the Control APR mean, and the other from the HEX APR mean. Removing these two women from the analysis, the effect in women is APR control = 1.07 ± 1.03, APR HEX = 1.32 ± 1.31, Wilcoxon signed rank test: Z = 2.15, p = 0.03, effect size r = 0.43. If we do not remove these two outliers, we get APR control = 1.34 ± 1.46, APR HEX = 1.66 ± 1.76, Wilcoxon signed rank test: Z = 1.90, p = 0.06, effect size r = 0.41] (Figure 4C). Thus, in two consecutive experiments, one across-participants and one within-participants, HEX decreased aggression in men, but increased aggression in women.

**Figure 4.**
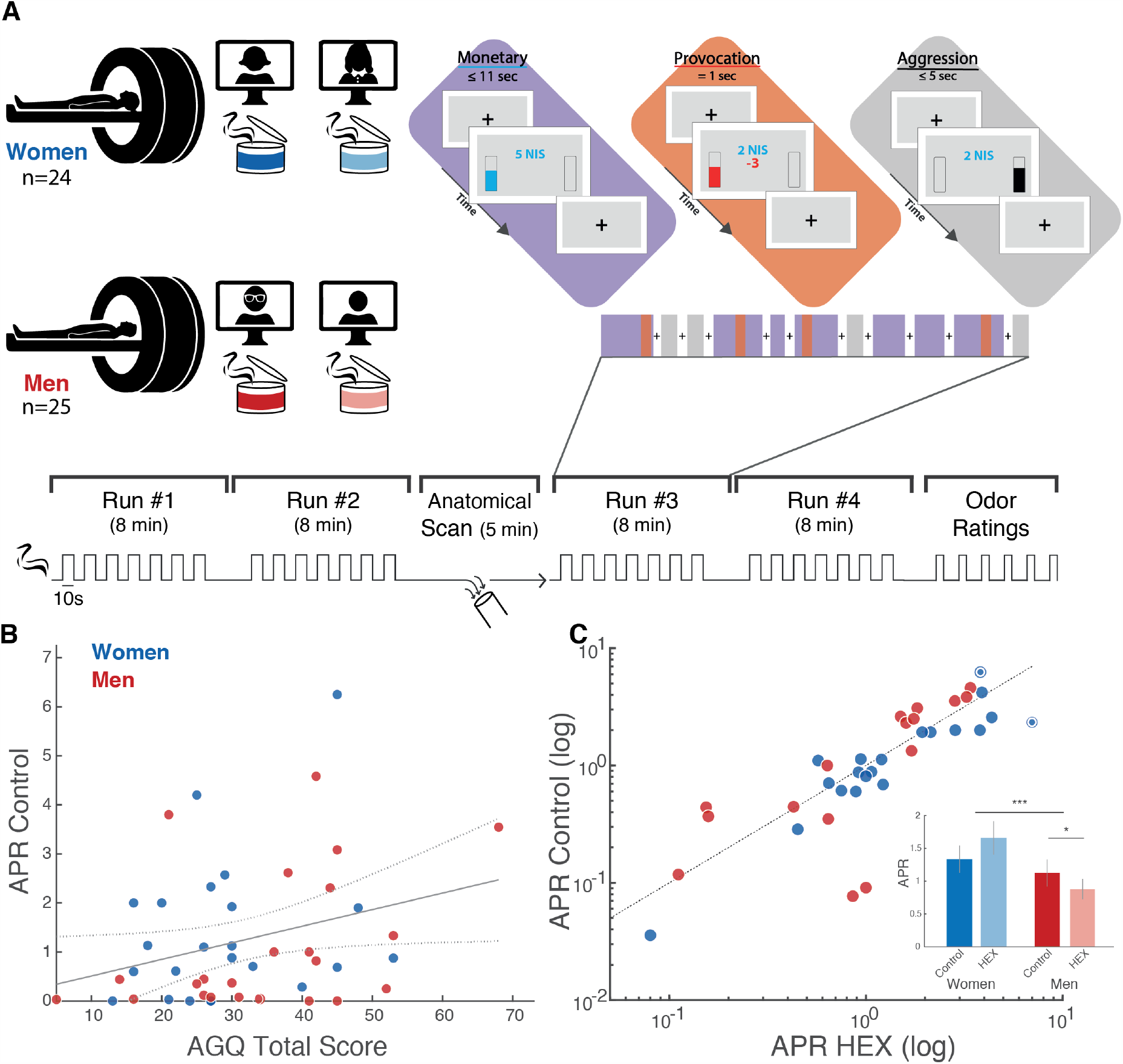
Exposure to HEX modulated aggressive behavior in a sex-dependent manner within participants. (**A**) In a within-subjects design, participants were exposed to an odorant (HEX and Control, counter-balanced for order), and played a game where their online partner stole their money occasionally (Provocation). During the game they could aggress against that same (non-existent) person by deducting money from them (Aggression). (**B**) Correlation between basal levels of aggression as measured here (aggression/provocation ratio - APR in the control condition) and an aggression questionnaire. (**C**) Scatter plot of the APR under HEX and the APR in the control condition. Each dot is a participant, women (blue) and men (red). The scatter plot is summarized in a bar graph. Dotted circles represent outliers (>3 s.d.). All tests were two-tailed, all centres reflect mean, all error-bars reflect SEM. * = p<0.05, *** = p<0.005.

### HEX increased activity in a brain area implicated in the perception of social cues

To investigate the impact of HEX on brain response, we conducted a whole-brain voxel-wise statistical parametric map. The ANOVA contrast of provocation vs. baseline (monetary) revealed a typical salience network activation (Figure 5).

In turn, the ANOVA contrast of provocation vs. baseline (monetary, see methods) with the added level of HEX vs. control in men and women revealed only one, but very pronounced significant activation in the left angular gyrus (AG), with no difference between men and women (Figure 5, Figure 6A). Although the statistical significance associated with this result is the mapping statistic (p < 0.001, corrected for multiple comparisons), in order to verify the directional drivers of this effect, we visualized the betas in the resultant left AG region of interest (Figure 6B, 6C). A two-tailed ANOVA applied to the beta values revealed a main effect of odour (F(97) = 6.79 p = 0.01), but no effect of sex (F(1,97) = 0, p = 0.95) and no interaction (F(1,97) = 0.03, p = 0.87). The main effect of odour reflected increased activation during provocation under HEX (mean beta (a.u.) HEX = 57.74 ± 22.76 (SE), mean beta (a.u.) control = −26.26 ± 22.37) (we reiterate that these statistics are merely to verify directionality, as the statistic associated with this finding remains the mapping statistic alone, namely p < 0.001, corrected for multiple comparisons). In previous research, the left AG has been implicated in the perception of social cues (*43*), and was recruited by perception of contextual social cues (*44*). Moreover, the left AG has been associated with brain mechanisms of aggression (*45*), and it is hypoactive in aggressive individuals (*46*). In other words, the AG is part of a social network related to aggression, and here it was recruited by a social putative chemosignal.

**Figure 5.**
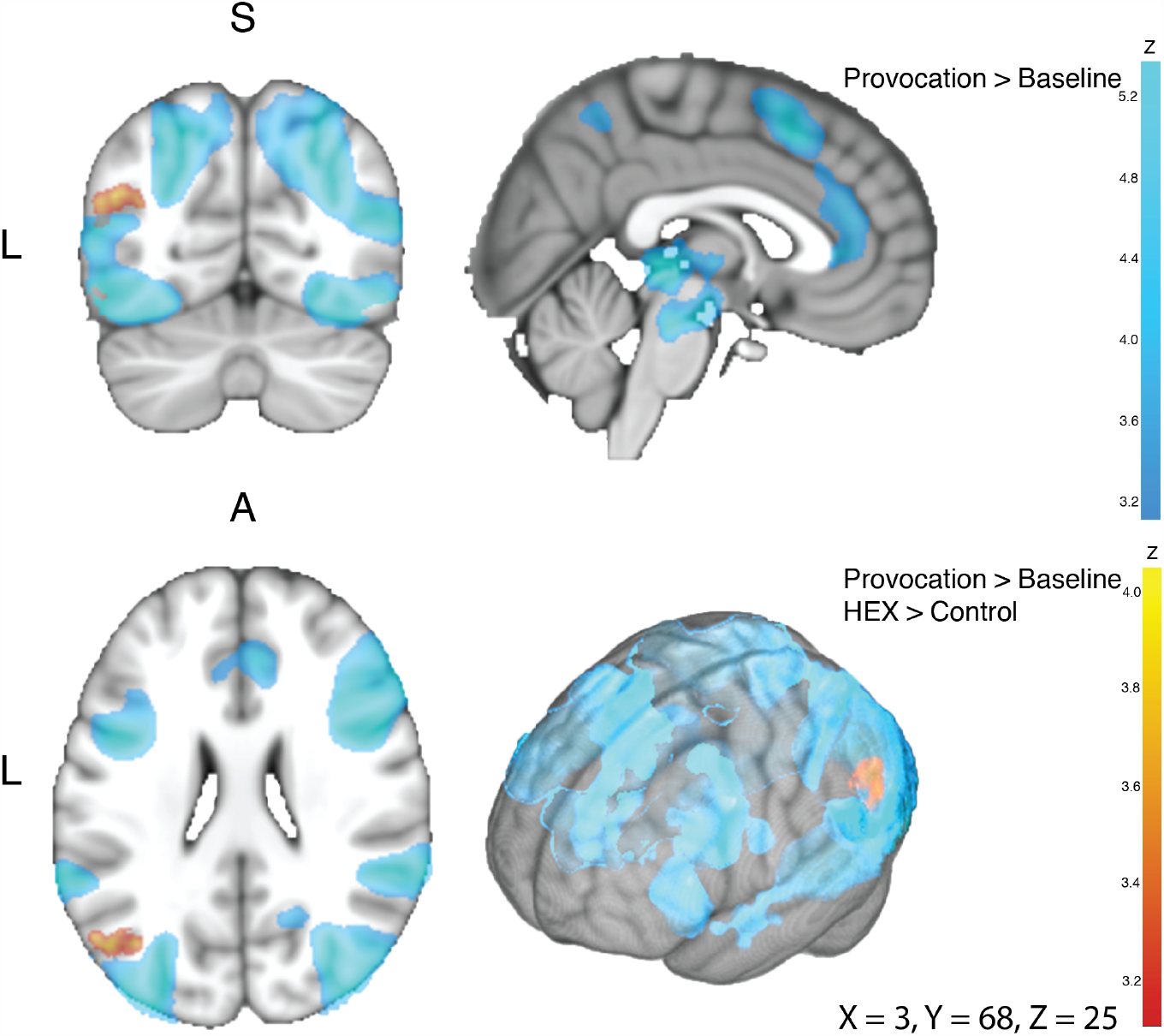
Provocation recruited an extensive brain network. The group image (n = 49), in coronal (top left), sagittal (top right), axial (bottom left) and surface (bottom right) views. In all panels, shades of blue reflect the Provocation > Baseline contrast, and shades of red reflect the Provocation > Baseline and HEX > Control contrast. In blue, provocation induced activity in the fusiform gyrus, OFC, Insula, Superior temporal gyrus, anterior cingulate cortex, inferior frontal gyrus, pre-SMA, precuneus, ventral tegmental area, periaqueductal grey area and thalamus (for full list, coordinates and peak activation, see Supplementary table 2). In red, HEX induced activation in left Angular gyrus.

### HEX modulated functional connectivity in brain networks of aggression

To investigate how the left AG may be modulating aggression under HEX, we investigated its functional connectivity with the entire brain. We applied whole-brain psychophysiological interaction (PPI) analysis (*47*) using the left functional AG region of interest (ROI) as a seed. We observed a pronounced dissociation by sex, that mirrored behavior. More specifically, HEX significantly modified left AG functional connectivity with three brain regions: The right temporal pole (extended to the middle temporal gyrus), the left amygdala (extended to the left hippocampus) and bilateral lateral orbitofrontal cortex (Figure 6D). Although the statistic associated with these HEX-related changes in functional connectivity is the PPI mapping statistic (p < 0.01 corrected for multiple comparisons [we note that the temporal pole result is stronger, surviving p < 0.001 corrected]), to further explore directionality, we applied ANOVAs to the beta values of connectivity. We observed that under provocation, HEX systematically increased connectivity to these regions in men, but decreased it in women. This effect recurred in the right temporal pole (F(1,97) = 5.60, p = 0.02, Men: mean beta-values HEX = 0.18 ± 0.16, mean beta-values Control = −0.25 ± 0.16. Women: mean beta-values HEX = −0.05 ± 0.16, mean beta-values Control = 0.20 ± 0.09. In the left amygdala (F(1,97) = 6.49, p-value = 0.01, Men: mean beta-values HEX = 0.14 ± 0.14, mean beta-values Control = −0.21 ± 0.11. Women: mean beta-values HEX = 0.01 ± 0.17, mean beta-values Control = 0.33 ± 0.09. And in the lateral OFC (F(1,97) = 8.40, p-value = 0.005, Men: mean beta-values HEX = 0.29 ± 0.21, mean beta-values Control = −0.31 ± 0.13. Women: mean beta-values HEX = −0.12 ± 0.24, mean beta-values Control = 0.33 ± 0.13 (Figure 6E, 6F) (we reiterate that these statistics are merely to verify directionality, as the statistic associated with this finding remains the mapping statistic alone, namely p < 0.01 corrected for multiple comparisons).

We observe that the temporal pole, amygdala, and OFC, are all parts of two brain-networks involved in aggressive behaviour; social and emotional evaluation and decision-making (*48*). Thus, our results imply that these regions may act in concert under the modulation of the AG, whereby increased functional connectivity with these regions is associated with reduced aggression (as in men), but decreased connectivity is associated with increased aggression (as in women). Finally, when looking at the shared change in FC for both women and men, we see an increase in FC for left motor areas: PMC and SMA (all participants were right handed, and used mostly the right hand for aggressive response). This is consistent with previous reports of increased activity in motor and pre-motor areas in response to provocations (*15*) (see Supplementary Figure 7).

## Discussion

Impulsive aggression is a major factor in the human condition, yet how exactly aggression is triggered or blocked in the human brain remains unclear (*15–17*). Moreover, real-world human impulsive aggression is one of the most sexually dimorphic behaviours (*49*), yet what brain mechanisms underlie this dimorphism also remains unclear (*17*). In animals ranging from insects to rodents, aggression is sexually dimorphic at levels ranging from genes to cells (*50, 51*), and this dimorphism has been linked to the olfactory system (*4, 52*). Here we find the same in humans. We observed that a body-volatile, namely HEX, significantly decreased aggression in men, yet significantly increased it in women. This same molecule also increased brain activity in the AG, a cross-modal integrating hub involved in social cognition (*53*). In other words, in humans, like in rodents, a “social odour” activates the “social brain”. Moreover, this same signal modulated functional connectivity between these substrates of social appraisal (AG) and a network previously associated with aggression. This included modulation of functional connectivity with the TP, an area similarly implicated in social appraisal (*54*) and aggression (*55*), and modulation of functional connectivity with the amygdala and OFC, namely substrates implicated in aggressive execution (*14, 16, 17*). All this occurred in a sex-dependent manner consistent with behaviour: HEX increased connectivity in men but decreased it in women. We would like to stress that in our analyses we did not first highlight the neural substrates of aggression and ask how they are impacted by HEX, but rather asked what brain networks are modulated by HEX, and this question happened to uncover the classic neural substrates of social appraisal and aggressive behaviour. We think this path of discovery significantly strengthens the idea that HEX is a human social chemosignal with relevance to aggression. Moreover, it provided for a behavioural and brain mechanistic depiction of human aggression and its sexual dimorphism: HEX may lead to increased or decreased aggression through increased or decreased control by the AG over the amygdala through a circuit involving the TP and OFC.

The sex dimorphism in our behavioural and brain results dovetails with previous findings obtained using functional brain imaging (*56*) to depict a level of functional brain sex-dimorphism in response to social odours that is not matched by any other sensory stimulus that we are aware of. Human functional brain responses to basic auditory and visual cues are generally non-dissociable by sex (*57*), yet here we could use them alone to discriminate men from women at 79.6% accuracy. This begs the question: what behavioural setting could underlie selection for a body-odour that increases aggression in women but decreases it in men? Or in other words, what could be the ecological relevance of these results? In this respect, we call attention to the setting of infant rearing. Parents across cultures are encouraged to sniff their babies (*58*), an action that activates brain reward circuits in women (*59*). Although an initial analysis of baby-head volatiles did not report on HEX (*60*), in-depth follow-up by the same authors using GCXGC-MS uncovered hexadecanal at about 20% the level of heptanal, the most abundant aldehyde component (Mamiko Ozaki, personal communication). Moreover, we also note in this respect that infant rearing is the one social setting where humans have extensive exposure to conspecific faeces (*61*), a rich source of HEX (*27*). Our results imply that sniffing babies may increase aggression in mothers, but decrease aggression in fathers. Whereas maternal aggression has a direct positive impact on offspring survival in the animal world (*62, 63*), paternal aggression has a negative impact on offspring survival (*64*). This is because maternal aggression is typically directed at intruders, yet paternal aggression, and more-so non-paternal male aggression, is often directed at the offspring themselves (*65–67*). If babies had a mechanism at their disposal that increased aggression in women but decreased it in men, this would likely increase their survival. Our results portray just such a mechanism.

Given all of the above, should we label HEX as a human pheromone? Sniffing human bodily secretions such as sweat and tears drives assorted behavioural and physiological effects (*11, 68*), and body-odours may reflect assorted emotional states (*69*) including aggression (*18, 19*), but the identity of specific molecular components involved in human social chemosignaling has remained elusive (*70*). Here we identify one such component, namely HEX, whose effects can be seen as consistent with those of a mammalian pheromone (*71*). Previously, the steroidal molecules estratetraenol (EST) and androstadienone (AND) had been proposed as human pheromones, yet this labeling was often rejected, primarily because EST and AND don’t clearly trigger or block behavior, nor do they have obvious ecological relevance (*72, 73*). Here HEX had a pronounced effect on behavior, and moreover, on the behavior of aggression, a domain dominated by pheromonal communication in most mammals (*7*). The notion of pheromonal communication was once considered dependent on a functional vomeronasal system, a system that humans may not possess (*74*). More recent views, however, blur this distinction, and highlight pheromonal communication through the main olfactory system as well (*75–79*). Despite these arguments for labelling HEX as a human pheromone, our results fall short of granting us that license. This is primarily because of one limitation: Although we demonstrated that the effects emerge following HEX and do not emerge following a control odour, and although the effects persisted despite a lack of percept associated with HEX, we nevertheless did not test against additional molecules also present in body-odour (e.g., the above-mentioned heptanal). In other words, it remains possible that the signal in question is “body-odour” (albeit subliminal), and not necessarily HEX alone. We submit that this does not take away from the fundamental findings we present, namely that a social chemosignal can trigger or block aggression, the pronounced sexual dimorphism in this cascade, and identification of the brain substrates involved, but it does raise the possibility that HEX is not unique in this, and that other body-volatiles may have similar effects. This issue remains to be explored, and currently prevents us from labelling HEX alone as a human pheromone. Beyond this, we would like to acknowledge several limitations: First, regarding our imaging results, we reiterate that correlation is not causation. We indeed identify a brain pattern associated with HEX, and it is tempting to suggest that this pattern is responsible for the observed effects, yet this can only be proven by experiments where the brain mechanism is perturbed, experiments that are very difficult to conduct in human participants. Second, we also acknowledge that our suggested ecological relevance in infant rearing was not directly tested in this study. Although we think it is a plausible hypothesis, it remains to be experimentally verified, and here serves only as an example of possible ecological relevance for our results.

Despite the above limitations, we conclude in stating that sniffing the body-odour-constituent HEX blocks aggression in men but triggers aggression in women. HEX may exert its effects by modulating functional connectivity between the brain substrates of social appraisal and the brain substrates of aggressive execution. This places chemosignaling at the mechanistic heart of human aggression, and poses but one added example to the rapidly growing body of evidence implicating social chemosignaling as a major, albeit mostly subconscious, power in human behaviour.

## Acknowledgments

This work was funded by an ERC AdG grant (SocioSmell 670798) awarded to Noam Sobel. We thank Macia Buades-Rotger, Ulrike Kramer, Katja Bertsch, Jeanette Mumford, Yaara Yeshurun and Anat Arzi for their valuable guidance.

## Author Contributions

N.S, H.B, J.S and E.M. conceptualized the idea. E.M. planned and created the experiments, acquired, analyzed, interpreted and visualized the data. D.A. acquired men data in the TAP. E.M and T.W. analyzed imaging data, D.H., A.W. and E.M created the pressure sensor apparatus. E.L. and L.G. handled the olfactometer. E.M., D.K. and R.Z. performed the ELISA. A.R. wrote code for the TAP. R.W., T.S., Y.E.S., S.A, L.R., N.R., A.R., L.G. and E.L. helped run participants. E.M. and N.S. wrote the manuscript.

## Methods

### Data availability

All data of this manuscript will be publically available on publication. All the behavioural data are already available on Mendeley at doi:10.17632/6nkh8j8733.1. All brain imaging data will be uploaded to OpenNeuro on acceptance.

### Code availability

All experiment and analysis codes will be uploaded to https://gitlab.com/worg_wis/eva-aggression

### Taylor Aggression Paradigm (TAP)

#### Participants

To estimate the number of needed participants we conducted a power analysis. We used the effect size obtained with aggression-related chemosignals (*18*) and using G*Power software (*80*) estimated, at alpha = 0.05 and 80% power, a necessary sample of at least 105 participants. Based on this, a total of 127 participants (67 men, mean age 25.48 ± 3.46, range 21-34) were recruited to a modified Taylor aggression paradigm (TAP) (*30–33*) after providing written informed consent to procedures that were approved by the Weizmann Institute Institutional Review Board (IRB). Participants were recruited using ads, and had no history of psychiatric drug use or any neurological or nasal conditions.

#### Odorant delivery

We used methods we have applied extensively in the past (*29, 81*). In an across-subjects double-blind design, half of the participants were exposed to carrier alone (10% eugenol, 100ul, diluted in Propylene Glycol) (control), and half to HEX masked in 10% eugenol (100ul, 0.083M). In brief, before commencement of the TAP, participants were first exposed to the odorant. Exposure started with 11 rating trials (inter-trial-interval = 25 seconds). On each trial the participant sniffed from an unmarked odorant-jar for 3 seconds, and then rated pleasantness, intensity and familiarity of the odorant along a visual-analogue scale. After the ratings session, an adhesive pad containing 30ul of the odorant was pasted onto the upper lip of the participant for continued exposure throughout the experiment. Participants and experimenters were of the same sex (three alternating men experimenters, four alternating women experimenters), and it was also noted to participants that their unseen game-partner is of the same sex.

#### Provocation phase

Provocation is a necessary catalyst for aggression in laboratory studies (*82*). To this end, participants were first antagonized towards a purported game-partner (Figure 1) in a form of the ultimatum game (*83*). In five rounds, participants were asked to distribute an amount of money (∼9$) between themselves and their fictitious game-partner. If they were to reach agreement, the distributed amount would be added to their earnings in the experiment. However, the interaction was rigged such that the fictitious game-partner refused any fair distribution, and agreed only to distributions where he/she received almost the entire amount (>8$). In order to maintain reliability, the time it took the purported game-partner to respond to offers was U-shaped, shorter for low and high amounts. The apparent egregious behaviour of the fictitious game-partner served as the aggression provoking mechanism.

#### Aggression discharge phase

After the provoking phase, participants engaged in a modified Taylor aggression paradigm. Participants were again misled to believe that they are playing against the same partner (from the provocation phase). This time, in a reaction-time task where they compete to identify a change in shape of a target. If they were faster to identify, they could blast their opponent with an unpleasant noise (noise-blast) at a volume of their choosing. If their game-partner was faster, they would endure a noise-blast induced by their game-partner. Determining which of the participants was faster was done using a responsive calculation. This assured that participants will induce noise-blasts 16 out of 27 rounds, yet with respect to their actual reaction times. Additionally, this served the purpose of, again, maintaining reliability of the experimental setup. Noise-blasts were induced using a button box with 6 buttons labelled with informative faces (modified from a pain scale for children (*84*) (Figure 2A). The faces portrayed the intensity of the noise-blast, from mildest (1) to most severe (6). Noise-blasts that participants heard were randomized in length (∼4 seconds) and volume (Mean = 90 db, coming from a speaker in front of them).

#### Questionnaires

Prior to any experimental manipulation, self-reported mood was measured using a commonly applied 17-item mood questionnaire (*85*) (Supplementary Figure 1). At the end of the experiment, we asked participants to rate, using a VAS, general questions probing their social evaluation and willingness to interact with their game-partner (Supplementary Figure 8; Supplementary Figure 9). Additionally, participants completed the autism quotient questionnaire (AQ) (*86*), State anxiety questionnaire (STAI) (*87*) and the Buss and Perry aggression questionnaire (AGQ) (*41*) (Supplementary Figure 6).

#### Statistical analysis

Odor ratings were standardized with respect to the entire sample mean and s.d. Odour ratings did not distribute normally (Pleasantness TAP: Shapiro-Wilk W= 0.98, p = 0.001, intensity TAP: W = 0.96, p = 0.002. Pleasantness FC-PSAP: W = 0.98, p = 0.001, intensity FC-PSAP: W = 0.75, p = 2.2*10^−6^). Therefore, we applied a linear mixed model with factors of Sex and Odor, and random effect of participant. Noise-blasts were standardized with respect to the entire sample mean and s.d. Noise-blast volume was entered into a repeated-measures ordinal logistic regression analysis (Cumulative Link Mixed Model fitted with the Laplace approximation) with factors of Odorant (HEX/Control) and Sex (men/women), and random effects of Participant. We further explored the data using the trial-by-trial mean for each group (16 separate noise-blast trials for each participant). In order to quantify this, we first conducted a Mann-Whitney test to compare between groups within trial. Then, an independent t-test on the trial-by-trial z-values. For further validation, we used a random permutation test. We shuffled the odour conditions of the real data and conducted a t-test on the shuffled z-distributions. We repeated the procedure 10,000 times and created a distribution of the t-statistics, resulting from statistical tests on the shuffled z-distributions. For reaction times, we first z-scored with respect to the entire group’s mean and s.d. RT’s did not distribute normally (Shapiro-Wilk: W = 0.49, p<2.2^−16^). Therefore, we applied a linear-mixed model with factors of Odour, Sex, Round, and participant as a random effect. For the exit questionnaire: Responses were z-scores to each Participant’s range. Self-reported mood was analysed using a linear-mixed model with factors of gender, odour, and question, and participant as a random effect.

#### Exclusions: Participants

Three participants were excluded from the TAP analysis due to technical faults of the acquisition system. Five participants did not answer the exit questionnaire, but were included in all other analyses. **Trials:** In the reaction time analysis we removed outliers of > 3 s.d. This resulted in removal of 99 trials out of 3456. This constitutes 2.86% of total trials, and 18.5% of trials in the one participant with most exclusions.

### Fist-Clench Point Subtraction Aggression Paradigm (FC-PSAP)

#### Participants

To estimate the number of needed participants we conducted a power analysis. We used the effect size obtained in the TAP to estimate group size using G*Power software. This implied that at alpha = 0.05 and 80% power we need to test a sample of 50 participants. Based on this, a total of 58 participants, of which 49 were included in final analyses (24 women, mean age 26.98 ± 3.92, range 19-36) participated in a modified Point Subtraction Aggression Paradigm (PSAP), after providing written informed consent to procedures that were approved by the Wolfson Hospital Helsinki Committee, and the Weizmann Institute Institutional Review Board (IRB). Participants were recruited using ads, and had no history of psychiatric drug use or any neurological or nasal conditions. All participants were right-handed.

#### FC-PSAP

Participants were told that their goal is to earn as much money as possible, and that we are interested in the dynamics of the interaction between them and the other participants. Participants actuated the PSAP using fist-clench (FC) activated pressure sensors (MR-compatible). The fist-clench devices were built in-house. They constituted a rubber ball in the hands of the participant, that was linked via ¼-inch Tygon tubing to a pressure sensor (all Sensors, 1INCH-Dx-4V-MINI). The sensors were powered at 5v (DAQ NI USB-6008), and the resultant signal was read using custom software written in MATLAB (code available at GitLab https://gitlab.com/worg_wis/eva-aggression) and LabChart7 software (ADInstruments, New South Wales, Australia). The software includes electrical noise removal, and calculation of a threshold. In the FC-PSAP, participants pressed both sensors simultaneously to earn money (monetary event), using one sensor, no matter which, they deducted money from the other participant (aggression). Provocation onsets were pre-randomized, and held constant across participants. The experimental timeline was such that the first MRI head scout was performed while participants calibrated the pressure sensors using alternating strong and weak presses, repeated three times. This served both the purpose of adapting the system to individual differences in pressure applied and also having the participants accustomed to the use of the FC pressure sensors (FCPS). Ensuing run duration was 8 minutes. After completing two runs with one odor condition, an anatomical scan was obtained. During the duration of the anatomical scan (5 minutes), air in the bore was high throughput vacuumed. Before starting the third run, participants were told that their (fictitious) game-partner was replaced, and now they are interacting with a new participant. This was done to minimize transferring of emotions, strategies etc. between odor conditions.

#### Odorant delivery in the MRI

Odorants were delivered using a computer-controlled air-dilution olfactometer of the type we have used extensively in the past (*81, 88*). Here we also introduced one modification whereby rather than using a nose-mask for delivery, we converted the entire head-coil into a controlled olfactory microenvironment. The head-coil was first enclosed with clear Teflon-coated Plexiglas. The olfactometer Teflon nosel was placed ∼10 cm anterior to the nose, and produced a constant flow of olfactometer air (1.5 lpm) that carried embedded 10 s pulses of odorant (HEX or control). A powerful 2-inch vacuum hose was connected to the back of the head coil, generating a constant directional-flow environment from the nosel in front of the nose, passed the nose, and to the back of the head-coil for removal. Nasal airflow was constantly precisely monitored using a nasal cannula linked to spirometer (ML141, ADInstruments), and instrumentation amplifier (Power-Lab 16SP, ADInstruments). Since Hexadecanal at this concentration and mineral oil have no perceivable odor, we validated precise olfactometer timing with a photoionization detector (PID) (RAE systems, Model ppbRAE 3000). In each experiment, there was a total of 4 runs, 2 with HEX (CAS# 629-80-1, 0.083 M, supplied by TCI and Cayman), and two with control (mineral oil alone, CAS# 8042-47-5, Sigma-Aldrich). Order of conditions was counterbalances, and the given condition was unknown to participants or experimenters. Only two out of 58 participants reported that they smelled any odor during the experiment. Odor ratings were completed inside the scanner, after completion of the FC-PSAP. At the end of the experiment participants completed an exit-questionnaire probing their thoughts, strategies and attitudes towards their game partners, a general demographics questionnaire, AGQ, AQ, and STAI.

#### MRI Data acquisition

MRI Scanning was performed on a 3 Tesla Siemens MAGNETOM Prisma scanner, using a 32-channel head coil. Functional data were collected using a T2*-weighted gradient-echo planar imaging sequence. In each run there were 240 repetitions comprising 56 slices, axial slices tilted 15° toward the AC–PC plane (TR=2000 ms, TE=30 ms, flip angle=75°, field of view 240×240 mm, matrix size 96×96, slice thickness of 2.5 mm with no gap, 2.5×2.5 mm^2^ in-plane resolution), covering whole brain. Anatomical images were acquired at 3D T1-weighted magnetization prepared rapid gradient-echo sequence at high resolution (1×1×1 mm^3^ voxel, TR=2300 ms, TE=2.98 ms, inversion time=900 ms, flip angle=9°).

#### fMRI data analysis

Neuroimaging data was analyzed using FSL (5.0.9), FEAT (v6.00), SPM12 (for PPI analysis) and MATLAB (R2017b, R2018b). FMRI data processing was carried out using FEAT (FMRI Expert Analysis Tool) Version 6.00, part of FSL (FMRIB’s Software Library, www.fmrib.ox.ac.uk/fsl). Registration of the functional data to the high-resolution structural image was carried out using the boundary-based registration algorithm (*89*). Registration of the high resolution structural to standard space images was carried out using FLIRT (*90, 91*) and was then further refined using FNIRT nonlinear registration (*92, 93*). The following pre-statistics processing was applied; motion correction using MCFLIRT (*91*); non-brain removal using BET (*94*); spatial smoothing using a Gaussian kernel of FWHM 6mm; grand-mean intensity normalisation of the entire 4D dataset by a single multiplicative factor; high-pass temporal filtering (Gaussian-weighted least-squares straight line fitting, with sigma = 62.5 sec). For each participant, a first-level general linear model included three regressors of: monetary (<11 s), provocation (=1 s) and aggressive events (<5 s), onset and offset were determined according to the visual stimuli presented to participants. Only offset times of monetary events dependent on participants’ behavior. Additionally, we regressed out failed events (‘none’ regressor), temporal derivatives, and modeled out single TR’s with excessive motion according to frame-wise displacement >0.9. The signal was convolved with double-gamma Hemodynamic response function (HRF). First, we modeled each run separately for each participant (4 in total), then combined the runs, adding the odor present for each run (two runs exposed to HEX, two to control, counterbalanced). Then we grouped the data, while adding a parametric modulation of participants’ sex. Time-series statistical analysis was carried out using FILM with local autocorrelation correction (*95*). The second-level analysis, which separated different odor condition runs and combined same odor runs, was carried out using a fixed effects model, by forcing the random effects variance to zero in FLAME (FMRIB’s Local Analysis of Mixed Effects) (*96*), (*97*), (*98*). The group-level analysis, which added participants’ sex and averaged across all group, was carried out using FLAME (FMRIB’s Local Analysis of Mixed Effects) stage 1 (*96*), (*97*), (*98*). Z (Gaussianised T/F) statistic images were thresholded using clusters determined by Z > 2.3 and a (corrected) cluster significance threshold of P = 0.05 (*99*). Analysis focused on provocation events for several reasons. First, Since the response in question is impulsive aggression, the moment of the provocation is the moment when the reactive spontaneous response happens, this is, by definition, the provocation event. Additionally, provocation events were well defined temporally, and most reproducible across participants. Finally, some participants did not have enough aggressive responses to allow analysis. Moreover, some participants had aggressive responses only in one odor condition and not the other, which severely skews the analysis. In the manuscript we report a contrast of Provocation > Baseline. Notably, due to the particular design in which provocation events occurred only during monetary events, even though technically the contrast is defined as provocation Vs. baseline, it is de facto a Provocation > Monetary contrast that we are estimating. We refer to it throughout the text as Provocation > Baseline. Due to this design, we also refrain from calculating the % change, and refer to betas as a measure of activation.

#### PPI analysis

To explore functional connectivity with the AG during provocation we conducted a whole-brain psychophysiological interactions analysis (PPI) (*100*), using the angular gyrus (AG) region of interest (ROI) as a seed region. The AG ROI was functionally defined from the GLM analysis of the group-level. We conducted a whole-brain PPI analysis using FSL, with the first regressor being the provocation events regressor (psychological regressor), the second was the time-course of the AG ROI (physiological regressor), and the third was the PPI regressor of the convoluted response (interaction regressor) generated using SPM12. Other regressors were the task regressors as in the GLM model (monetary, aggression, none and motion according to frame-wise displacement>0.9). The following pre-statistics processing was applied; grand-mean intensity normalization of the entire 4D dataset by a single multiplicative factor. Time-series statistical analysis was carried out using FILM (*95*). Second-level analysis, in which we contrasted the odor conditions and combined same-odor runs, was carried out using a fixed effects model, by forcing the random effects variance to zero in FLAME (FMRIB’s Local Analysis of Mixed Effects) (*96*), (*97*), (*98*). Group-level analysis, in which we added the sex parameter and averaged across participants, was carried out using FLAME (FMRIB’s Local Analysis of Mixed Effects) stage 1 (*96*), (*97*), (*98*). Z (Gaussianised T/F) statistic images were thresholded using clusters determined by Z>2.58 and a (corrected) cluster significance threshold of P=0.05 (*99*).

#### Exclusions

One participant reported she did not believe the purported participant was real, and was therefore excluded from further analyses. Additional participants were excluded from analyses due to technical problems (n=5), AQ score above 32 (n=1 women) (in accordance with (*86*)), or excessive head-movements during the scan (n=1), retaining 24 women and 25 men in all reported analyses.

#### Data analysis tools

All analyses and statistical analyses were done using MATLAB R2018a and R2018b (The MathWorks, Inc.), FSL 5.0.9 (*101*), FEAT 6.00 (*95, 97*), SPM12 (*102*), and R 3.6.1 (*103*) (packages: readxl version 1.3.1, tidyverse 1.2.1, Hmisc version 4.3-0, plyr version 1.8.4, RColorBrewer version 1.1-2, reshape2, lme4 version 1.1-21, emmeans version 1.3.5, nlme version 3.1-140, pwr version 1.3-0, DescTools version 0.99.35).

