## Supplementary materials (Figures and Tables) for "Sniffing the Human Body-Volatile Hexadecanal Blocks Aggression in Men but Triggers Aggression in Women"

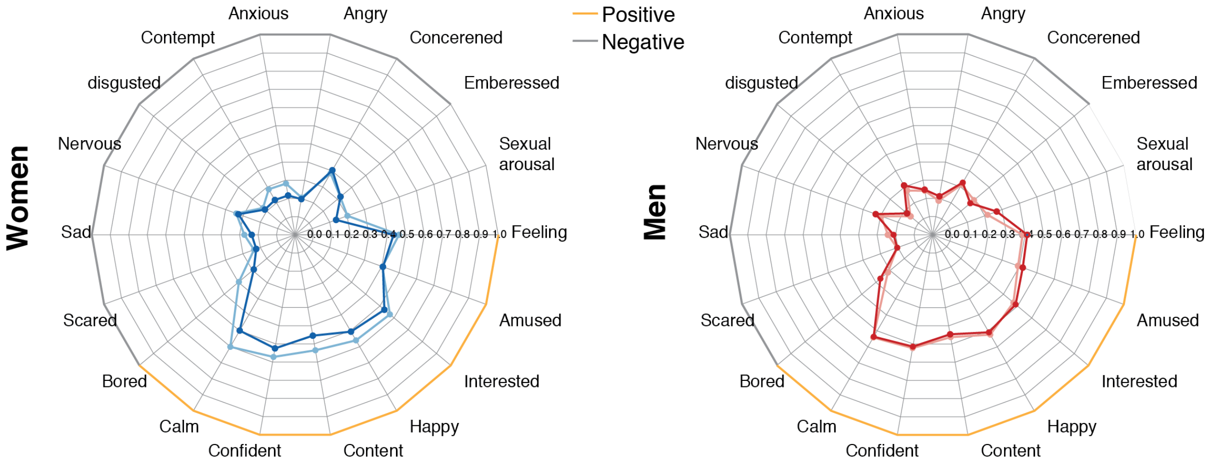

**Supplementary Figure 1. Participants' mood did not differ between groups prior to odor exposure** Participants' mood prior to experiment was estimated using a 17-item mood scale. A linear mixed-model with factors of Sex, Odor and Question, and random effects of participant revealed an effect of Question ($F(17) = 146.06$, $p < 0.0001$), and a modest significant interaction of Sex x Question ($F(17) = 1.74$, $p = 0.03$) indicating no initial differences within Sex groups.

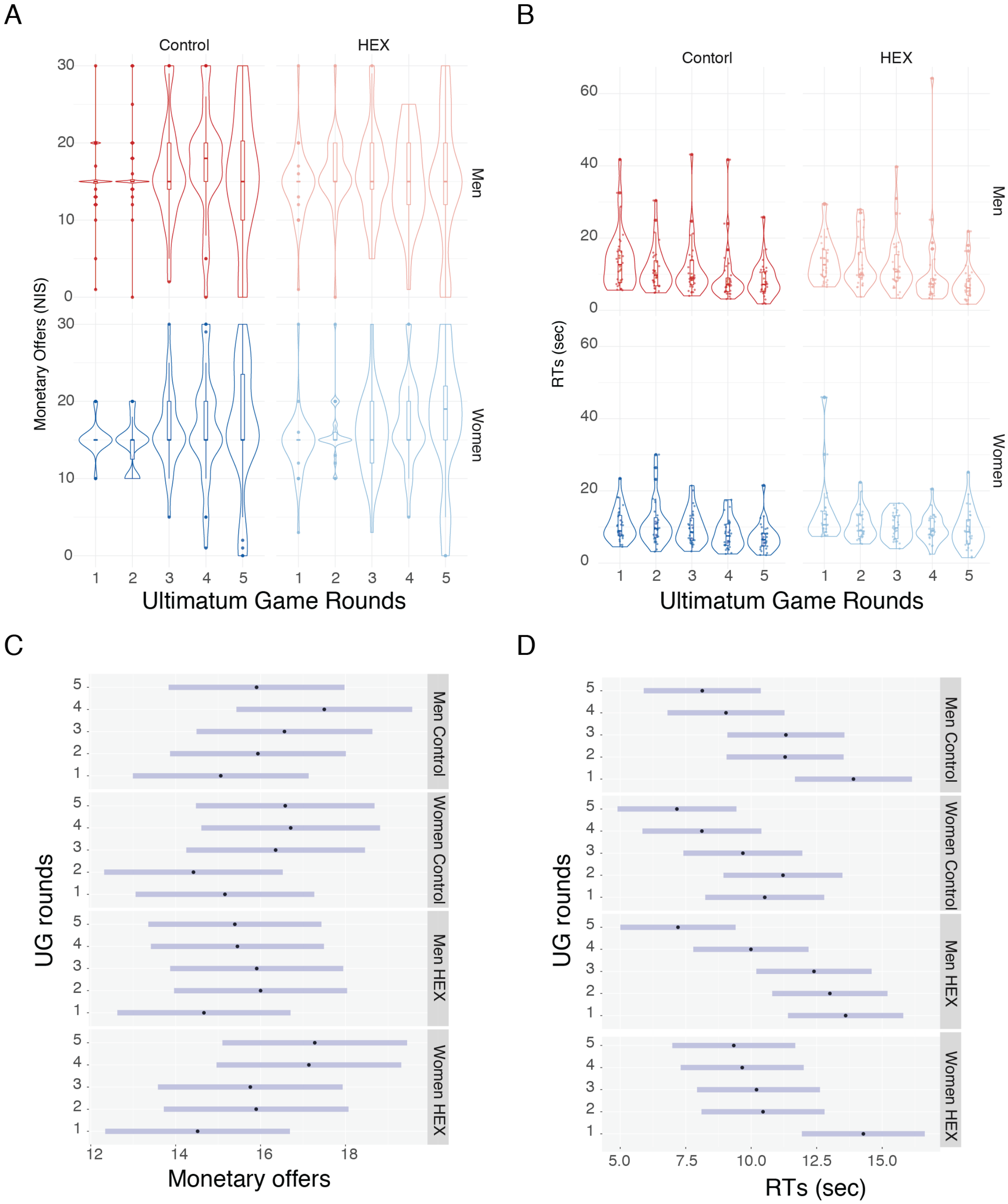

**Supplementary Figure 2. Participants' offers in the ultimatum game did not differ between groups**

(**A**) Participants' monetary offers were compared using a linear mixed-model with factors of Odor, Sex, Round and random effects of participant. There were no significant differences between groups (Sex: F(1) = 0.048, p = 0.83) , Odor: F(1) = 0.13, p = 0.72), Round: F(4) = 2.32, p = 0.06), Sex x Odor: F(1) = 0.56, p = 0.46, Sex x Round: F(1) = 0.71, p = 0.59, Odor x Round: F(1) = 0.71, p = 0.59, Odor x Round x Sex: F(1) = 0.71, p = 0.59). (**B**) Linear mixed-model with factors of Odor, Sex, Rounds, and random effects of participant revealed no significant effect of Odor and Sex on RT normalized score (Sex: $F(1) = 1.52$, $p = 0.22$, Odor: $F(1) = 1.52$, $p = 0.22$, Sex x Odor: $F(1) = 0.37$, $p = 0.54$, Sex x Rounds: $F(4) = 1.04$, $p = 0.39$, Odor x Rounds: $F(4) = 0.26$, $p = 0.90$, Sex x Odor x Rounds: $F(4) = 2.01$, $p = 0.09$) and a significant effect of Round (Rounds: $F(4) = 18.37$, $p < 0.0001$ ), which looks like a drift towards shorter RTs. This may imply that exposure to HEX did not reduce responsiveness or dampened participants' ability. The differences between the group in both monetary offers and RT's are better visualized using the (**C**) Mean + confidence intervals (CI) of monetary offers, and (**D**) Mean + CI of RTs.

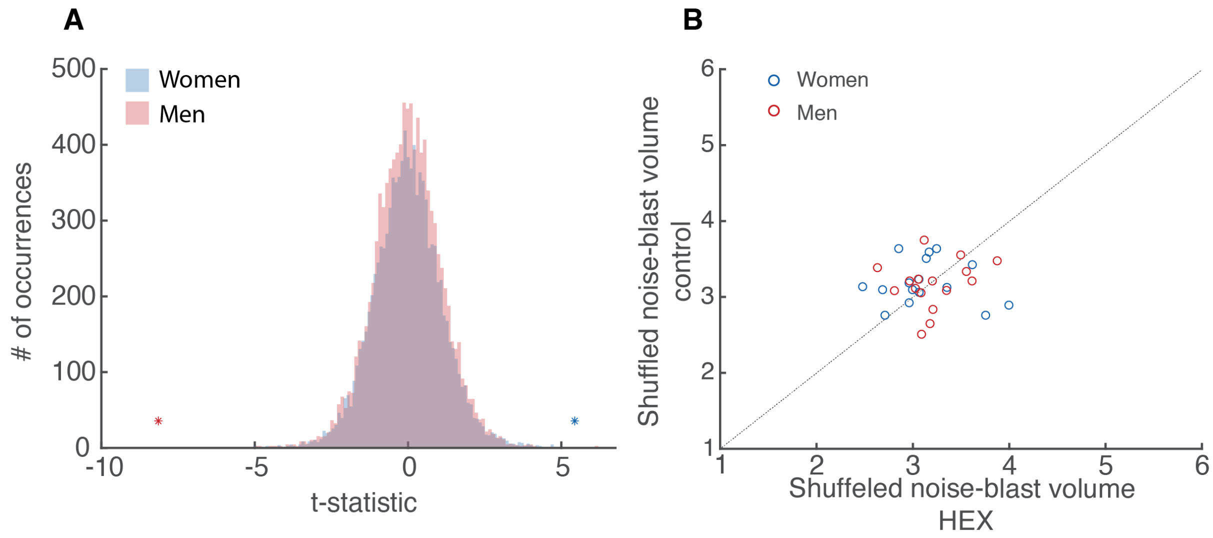

**Supplementary Figure 3. Validation of group differences using a permutation test**
(**A**) Distribution of t-statistics of shuffled data trial-by-trial comparisons. Asterisks denote the real data p-value, which is smaller than 99%. (**B**) Rendition of Fig. 2B using permuted data (see methods, Taylor Aggression Paradigm (TAP), Statistical analysis).

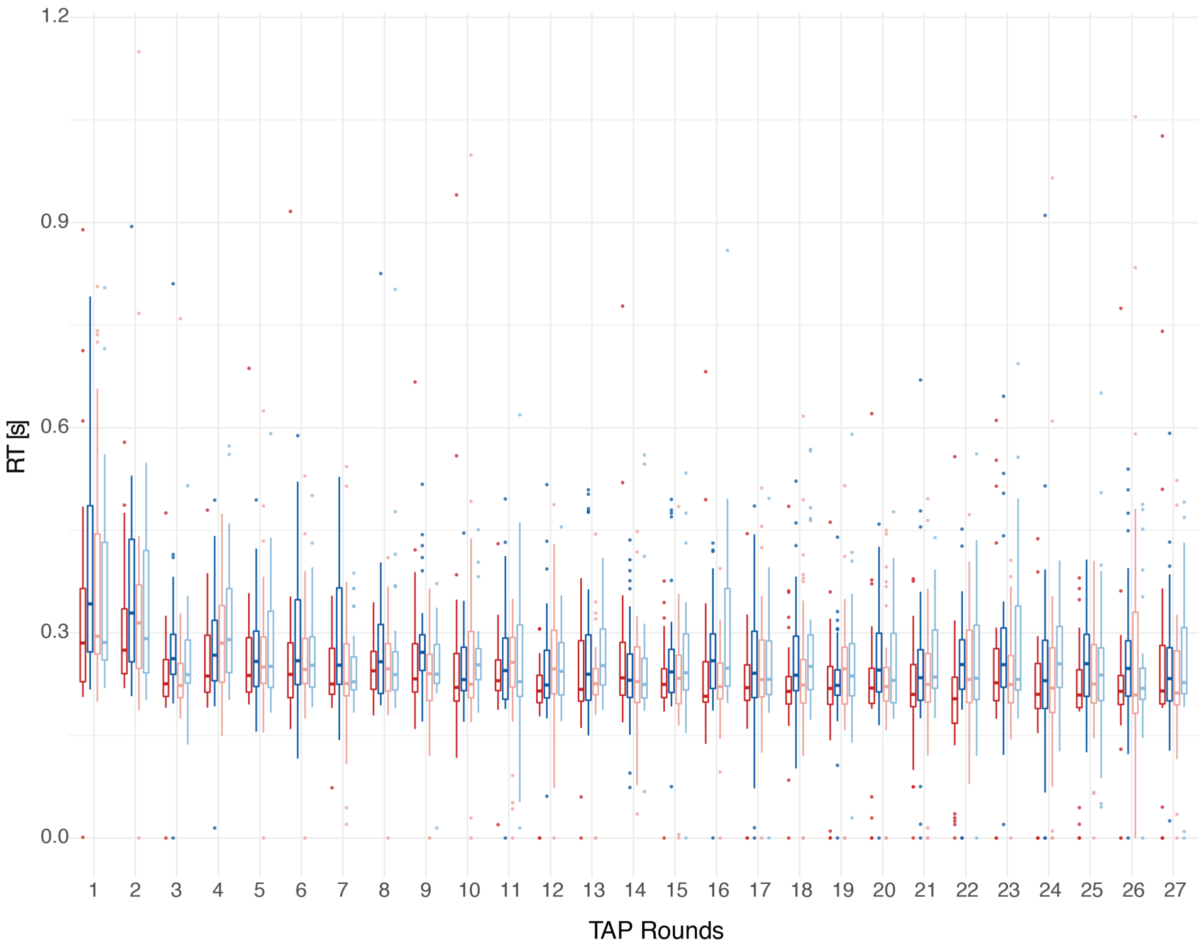

**Supplementary Figure 4. HEX does not affect vigilance in the TAP**

Linear-mixed model with factor of Odor, Sex, Round and random effects of Participant revealed a significant difference between Rounds, but no other significant difference (Sex: $F(1) = 0.02$, $p = 0.87$, Odor: $F(1) = 0.38$, $p = 0.54$, Round: $F(26) = 5.60$, $p < 0.0001$, Sex x Odor: $F(1) = 0.20$, $p = 0.65$, Sex x Rounds: $F(26) = 0.30$, $p = 0.99$, Odor x Rounds: $F(26) = 1.10$, $p = 0.33$, Sex x Odor x Rounds: $F(26) = 0.79$, $p = 0.77$). This indicated that there was no motor/cognitive difference between the groups. Boxplot depicts 25-75 percentile, line depicts median, bars denote min and max, dots depict data point larger than 1.5 s.d. in the data.

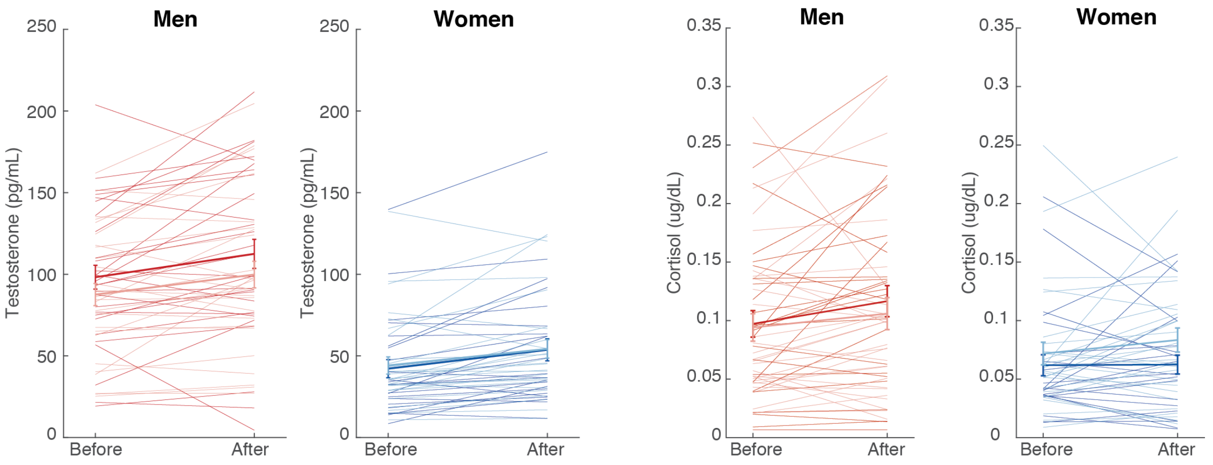

**Supplementary Figure 5. Salivary Testosterone and Cortisol levels did not differ between conditions**

Salivary Cortisol (CORT) and Testosterone (T) were measured prior and after the experiment. Saliva was collected before exposure to odor and at the end of the experiment. Participants rinsed their mouth with water before first saliva collection and passively drooled into a salivette. Samples were stored at -20C$^0$ and thawed and centrifuged prior to analysis using cortisol and testosterone kits (Cortisol Immunoassay kit, Testosterone Immunoassay kit, Salimetrics, CA, USA). After completion of the immunoassay, the absorbance of the fluorescent cortisol conjugate–antibody complex in the wells were obtained at 450 nm and corrected at 490 nm with a microplate reader. Standard dilutions of cortisol (0, 0.012, 0.037, 0.111, 0.333, 1.0 and 3.0 µg/dL) and testosterone (0, 6.1, 15.4, 38.4, 96, 240 and 600 pg/mL) were used along a nonspecific binding well in the first two columns of the kit for calibration. Defined high and low control concentrations were used as quality controls for each column of the plate. The absolute salivary cortisol and testosterone concentrations were estimated from the fluorescence of the hormone conjugate–antibody complex by computing the inverse value on a four-parameter sigmoid fit obtained with the standard values. For each subject, the delta of the of the two samples was computed.

An ANOVA on the ratio of before and after exposure to odor with a factor of Sex and Odor revealed a marginal difference between men and women's initial T levels ($F(1) = 3.7$ $p = 0.06$), but no effect of Odor levels ($F(1) = 0.98$ $p = 0.32$) and no interaction ($F(1) = 1.11$ $p = 0.30$) in T levels. An ANOVA applied on the CORT ratio revealed no effect of Odor ($F(1) = 0.18$ $p = 0.67$), Sex ($F(1) = 0.21$ $p = 0.65$), nor an interaction ($F(1) = 0.35$ $p = 0.87$). This might be due to the small group sizes for this kind of between-subject effects, given the inherent noise in measuring hormone levels.

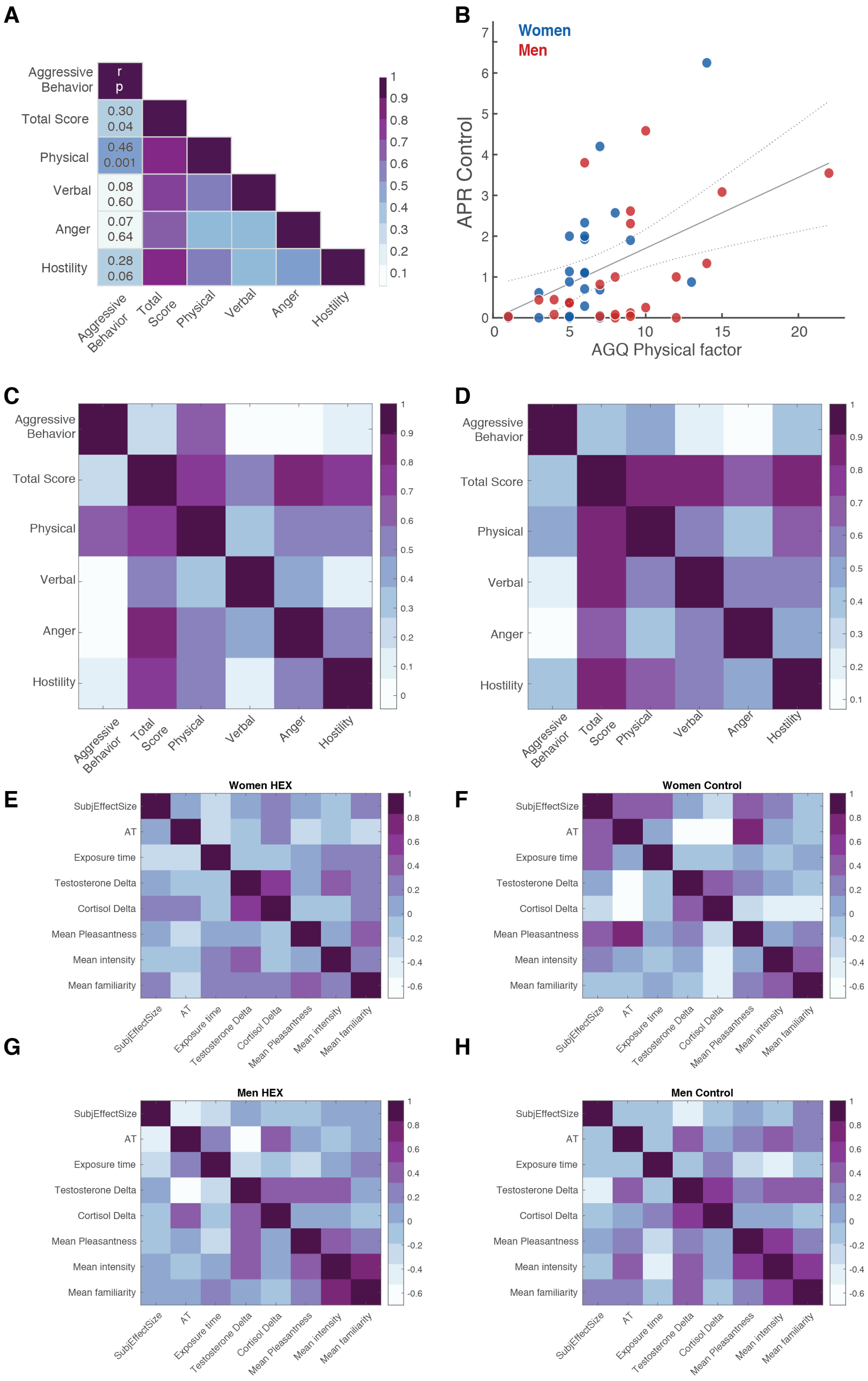
A
Aggressive Behavior
Total Score
Physical
Verbal
Anger
Hostility
r
p
0.30
0.04
0.46
0.001
0.08
0.60
0.07
0.64
0.28
0.06
B
Women
Men
APR Control
AGQ Physical factor
C
D
E
Women HEX
F
Women Control
G
Men HEX
H
Men Control
SubjEffectSize
AT
Exposure time
Testosterone Delta
Cortisol Delta
Mean Pleasantness
Mean intensity
Mean familiarity

**Supplementary Figure 6. FC-PSAP captures trait aggression as measured by the AGQ**

(**A**) Correlation matrix of AGQ; total score and the different factors with APR in control condition. (**B**) Correlation of the physical aggression factor in the AGQ with APR in control condition. r = 0.47, p = 0.001. The physical aggression factor usually correlated better than the total score of the AGQ in laboratory paradigms of aggression. The same correlation matrix, separated for women (**C**) and men (**D**), (**E**), (**F**), (**G**), (**H**) Subject effect sizes (computed for all subjects using the subject mean minus group mean in control divided by the pooled variance of the control and HEX groups) do not correlate with intensity nor with pleasantness ratings.

| Subject | Sex | Agg HEX | Agg Control | Mon HEX | Mon Ctrl | Prov HEX | Prov Ctrl | None HEX | None Ctrl |
|---|---|---|---|---|---|---|---|---|---|
| 1 | w | 32 | 42 | 35 | 32 | 9 | 10 | 6 | 5 |
| 2 | m | 0 | 4 | 54 | 49 | 17 | 15 | 4 | 6 |
| 3 | m | 0 | 0 | 58 | 58 | 18 | 20 | 0 | 1 |
| 4 | m | 6 | 8 | 52 | 59 | 14 | 18 | 6 | 1 |
| 5 | m | 9 | 7 | 51 | 54 | 14 | 20 | 5 | 1 |
| 6 | m | 2 | 2 | 55 | 57 | 18 | 17 | 4 | 0 |
| 7 | m | 3 | 7 | 54 | 51 | 19 | 19 | 4 | 4 |
| 8 | w | 1 | 0 | 56 | 56 | 20 | 18 | 1 | 3 |
| 9 | m | 16 | 30 | 43 | 41 | 10 | 13 | 8 | 3 |
| 10 | w | 11 | 12 | 55 | 50 | 17 | 17 | 0 | 3 |
| 11 | w | 16 | 15 | 50 | 53 | 15 | 17 | 1 | 0 |
| 12 | w | 5 | 0 | 54 | 58 | 17 | 17 | 2 | 1 |
| 13 | w | 16 | 17 | 49 | 49 | 17 | 15 | 0 | 0 |
| 14 | m | 31 | 37 | 42 | 37 | 17 | 12 | 4 | 1 |
| 15 | m | 24 | 34 | 47 | 41 | 16 | 13 | 0 | 0 |
| 16 | m | 34 | 39 | 44 | 41 | 12 | 11 | 3 | 0 |
| 17 | m | 18 | 2 | 48 | 58 | 21 | 26 | 0 | 0 |
| 18 | w | 15 | 11 | 45 | 46 | 20 | 18 | 7 | 12 |
| 19 | w | 18 | 18 | 46 | 40 | 15 | 16 | 6 | 13 |
| 20 | w | 0 | 0 | 58 | 57 | 26 | 28 | 0 | 2 |
| 21 | w | 0 | 0 | 57 | 54 | 25 | 23 | 3 | 6 |
| 22 | w | 11 | 7 | 69 | 73 | 12 | 8 | 2 | 1 |
| 23 | m | 2 | 0 | 57 | 56 | 24 | 28 | 1 | 4 |
| 24 | w | 48 | 36 | 32 | 41 | 11 | 14 | 0 | 2 |
| 25 | m | 29 | 24 | 42 | 45 | 17 | 18 | 0 | 0 |
| 26 | m | 0 | 3 | 69 | 62 | 10 | 21 | 9 | 0 |
| 27 | m | 0 | 18 | 57 | 48 | 26 | 22 | 0 | 0 |
| 28 | w | 32 | 25 | 40 | 50 | 15 | 13 | 3 | 2 |
| 29 | m | 0 | 2 | 58 | 58 | 27 | 26 | 1 | 0 |
| 30 | w | 12 | 6 | 57 | 72 | 20 | 10 | 0 | 0 |
| 31 | w | 12 | 26 | 49 | 40 | 21 | 20 | 0 | 0 |
| 32 | m | 0 | 1 | 56 | 56 | 27 | 25 | 3 | 3 |
| 33 | w | 9 | 4 | 52 | 58 | 20 | 14 | 4 | 11 |
| 34 | m | 21 | 15 | 33 | 26 | 12 | 6 | 6 | 0 |
| 35 | w | 22 | 11 | 46 | 61 | 18 | 16 | 1 | 1 |
| 36 | m | 34 | 55 | 25 | 29 | 10 | 12 | 22 | 0 |
| 37 | m | 4 | 11 | 56 | 53 | 26 | 25 | 0 | 0 |

| | | | | | | | | | |
|---|---|---|---|---|---|---|---|---|---|
| 38 | m | 14 | 22 | 52 | 45 | 22 | 22 | 0 | 0 |
| 39 | w | 37 | 30 | 37 | 38 | 13 | 15 | 0 | 5 |
| 40 | m | 0 | 1 | 57 | 58 | 26 | 27 | 1 | 0 |
| 41 | m | 21 | 22 | 50 | 49 | 20 | 22 | 0 | 1 |
| 42 | m | 0 | 1 | 57 | 57 | 27 | 26 | 2 | 1 |
| 43 | w | 49 | 35 | 32 | 37 | 7 | 15 | 2 | 3 |
| 44 | w | 2 | 1 | 57 | 56 | 25 | 28 | 0 | 2 |
| 45 | w | 38 | 28 | 34 | 41 | 10 | 14 | 3 | 5 |
| 46 | w | 42 | 50 | 34 | 28 | 11 | 8 | 0 | 1 |
| 47 | w | 13 | 17 | 39 | 51 | 13 | 21 | 25 | 4 |
| 48 | w | 29 | 25 | 40 | 52 | 15 | 13 | 2 | 0 |
| 49 | m | 26 | 23 | 50 | 61 | 8 | 6 | 5 | 0 |

**Supplementary Table 1. Number of events for each participant under HEX and under Control**

Events were coded as None in the case that participants had a technical problem or that they chose not to respond. Mean number of aggressive events under HEX condition: M = 15.59±14.28, Mean number of aggressive events under Control condition: M = 16.00±14.71, Mean number of Monetary events under HEX M = 48.76±9.66, Mean number of Monetary events under Control condition: M = 49.84±10.40, Provocation HEX M = 17.51±5.94, Mean number of None events under HEX condition: M = 3.18±4.90, Mean number of None events under Control condition: M = 2.20±3.10.

| Provocation > Baseline | | | | |
|---|---|---|---|---|
| **Z** | **Peak activation coordinates** | | | **Area** |
| | **x** | **y** | **z** | |
| **8.75** | -36 | -52 | -17 | Fusiform, left |
| **8.09** | -42 | -58 | -10 | Occipital lobe, left |
| **8.02** | 38 | -56 | -13 | Fusiform, right |
| **7.92** | 37 | -84 | -8 | Occipital lobe, right |
| **7.56** | 44 | 12 | 28 | Inferior frontal gyrus, right |
| 7.28 | 33 | -54 | 47 | Parietal cortex, right |
| **6.89** | 29 | 19 | -15 | Orbitofrontal cortex, right |
| **6.88** | -34 | 18 | -15 | Orbitofrontal cortex, left |
| **6.86** | -32 | -58 | 48 | Parietal cortex, left |
| **6.42** | -2 | -29 | -3 | Periaqueductal gray, left |
| **6.33** | -42 | 3 | 26 | Inferior frontal gyrus, left |
| **6.3** | 2 | -28 | -2 | Periaqueductal gray, right |
| **5.74** | 6 | 17 | 49 | Pre SMA, right |
| **5.5** | 2 | -26 | -23 | Ventral tegmental area, right |
| **5.25** | 53 | 4 | -14 | Superior temporal gyrus, right |
| **5.2** | -32 | 19 | 3 | Insula, left |
| **5.12** | 33 | 23 | 3 | Insula, right |
| **5.08** | -1 | -27 | -23 | Ventral tegmental area, left |
| **4.63** | 44 | 21 | -25 | Temporal pole, right |
| **4.6** | 6 | 34 | 23 | ACC, right |
| **4.46** | 6 | -8 | 1 | Thalamus, right |
| 4.3 | -7 | -8 | -1 | Thalamus, left |
| **4.26** | -6 | 17 | 43 | Pre SMA, left |
| **4.05** | -44 | 6 | -20 | Temporal pole, left |
| **3.85** | 7 | -59 | 53 | Precuneus, right |
| 3.8 | -6 | 34 | 17 | ACC, left |
| **3.57** | -51 | 1 | -14 | Superior temporal gyrus, left |
| 3.43 | -8 | -59 | 51 | Precuneus, left |
| 3.99 | -48 | -70 | 26 | Angular gyrus, left |

**Supplementary Table 2. Brain areas activated by provocation events compared to baseline** Coordinates and Zstat for all significant activation (cluster-corrected >3.1) for the contrast Provocation > Baseline.

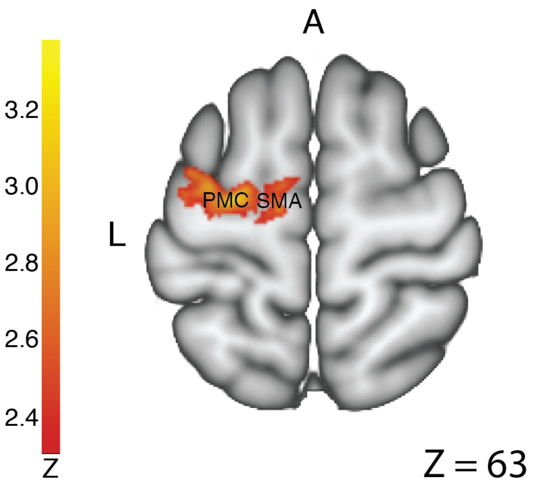

**Supplementary Figure 7. Pre-motor and supplementary motor areas functional connectivity with the AG was increased upon provocation when exposed to HEX**

PPI analysis with the AG as seed region revealed that the premotor cortex and supplementary motor area were more connected to the AG in both women and men for the contrast HEX>Control, Provocation>Baseline.

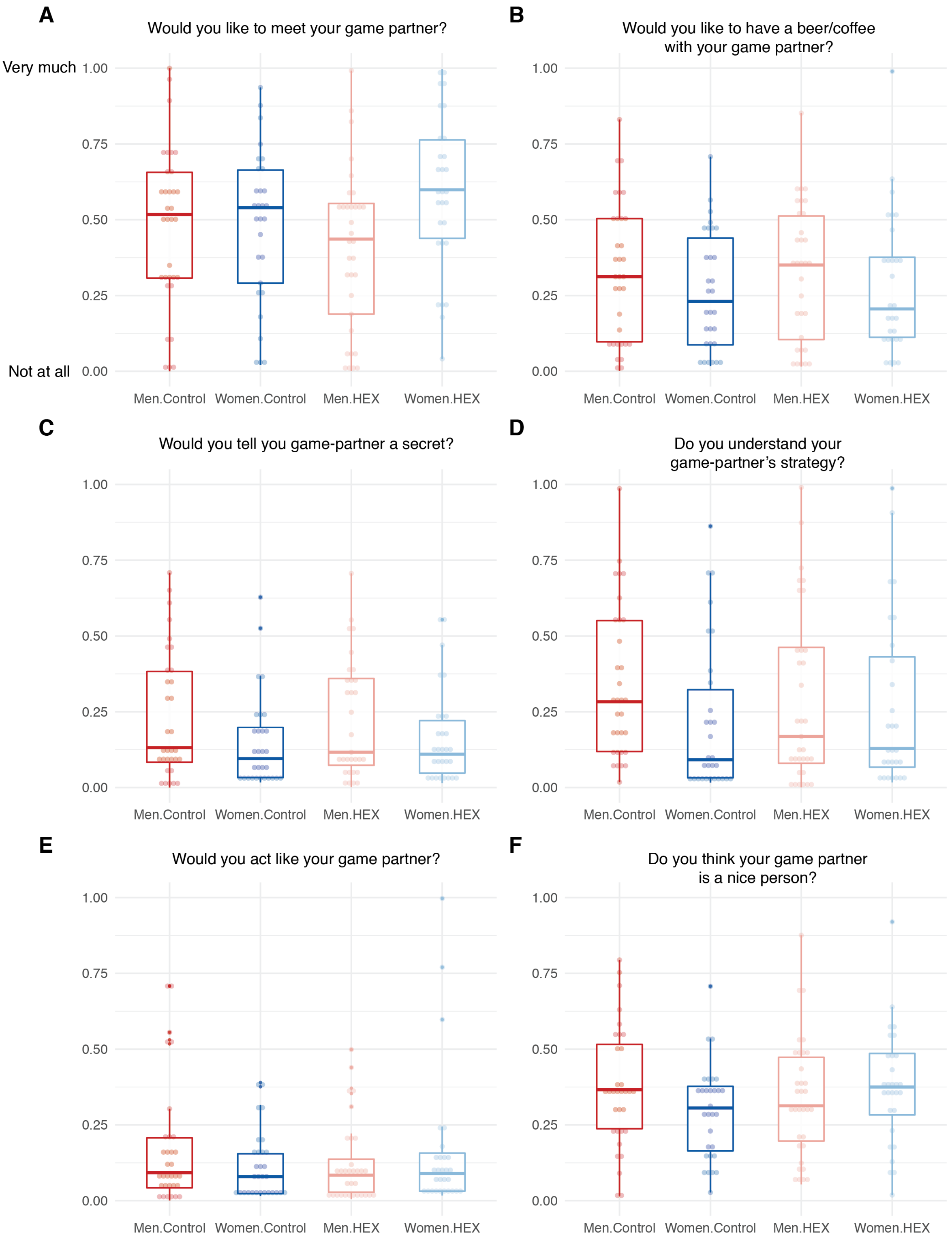
A
Would you like to meet your game partner?
Very much
Not at all
B
Would you like to have a beer/coffee with your game partner?
C
Would you tell you game-partner a secret?
D
Do you understand your game-partner's strategy?
E
Would you act like your game partner?
F
Do you think your game partner is a nice person?
1.00
0.75
0.50
0.25
0.00
Men.Control
Women.Control
Men.HEX
Women.HEX

**Supplementary Fig. 8. Participants' attitudes towards their game-partner during the TAP**

(**A**) *Would you like to meet your game-partner?* Linear mixed model with factors of Odor, Sex and random effects of participant revealed a main effect of Sex ($F(1) = 4.92$, $p = 0.03$), no effect of Odor ($F(1) = 0.0009$, $p = 0.98$) and a trend for an Sex x Odor effect ($F(1) = 3.20$, $p = 0.08$). Post-hoc analysis of the pairwise comparison revealed a significant difference between women and men in the HEX condition ($t(121) = 2.83$, $p = 0.03$). (**B**) *Would you like to have a beer or coffee with your game-partner?* Linear mixed model with factors of Odor, Sex and random effects of participant revealed no effects (Sex: $F(1) = 1.40$, $p = 0.24$, Odor: $F(1) = 0.03$, $p = 0.86$, Sex x Odor: $F(1) = 0.12$, $p = 0.73$). (**C**) *Would you tell your game-partner a secret?* Linear mixed model with factors of Odor, Sex and random effects of participant revealed a marginal effect of (Sex: $F(1) = 3.79$, $p = 0.05$, no effect of Odor: $F(1) = 0.0004$, $p = 0.98$, and no interaction Sex x Odor: $F(1) = 0.40$, $p = 0.53$). (**D**) *Do you understand your game-partner's strategy?* Linear mixed model with factors of Odor, Sex and random effects of participant revealed no effects (Sex: $F(1) = 1.34$, $p = 0.25$, Odor: $F(1) = 0.08$, $p = 0.77$, Sex x Odor: $F(1) = 1.34$, $p = 0.25$) (**E**) *Would you act like your game partner did?* Linear mixed model with factors of Odor, Sex and random effects of participant revealed no effects (Sex: $F(1) = 0.07$, $p = 0.79$, Odor: $F(1) = 0.04$, $p = 0.85$, Sex x Odor: $F(1) = 2.69$, $p = 0.10$). (**F**) *Do you think your game-partner is a nice person?* Linear mixed model with factors of Odor, Sex and random effects of participant revealed no main effect of Sex ($F(1) = 0.37$, $p = 0.54$), no effect of Odor ($F(1) = 0.42$, $p = 0.52$) and a trend for an Sex x Odor effect ($F(1) = 3.05$, $p = 0.08$).

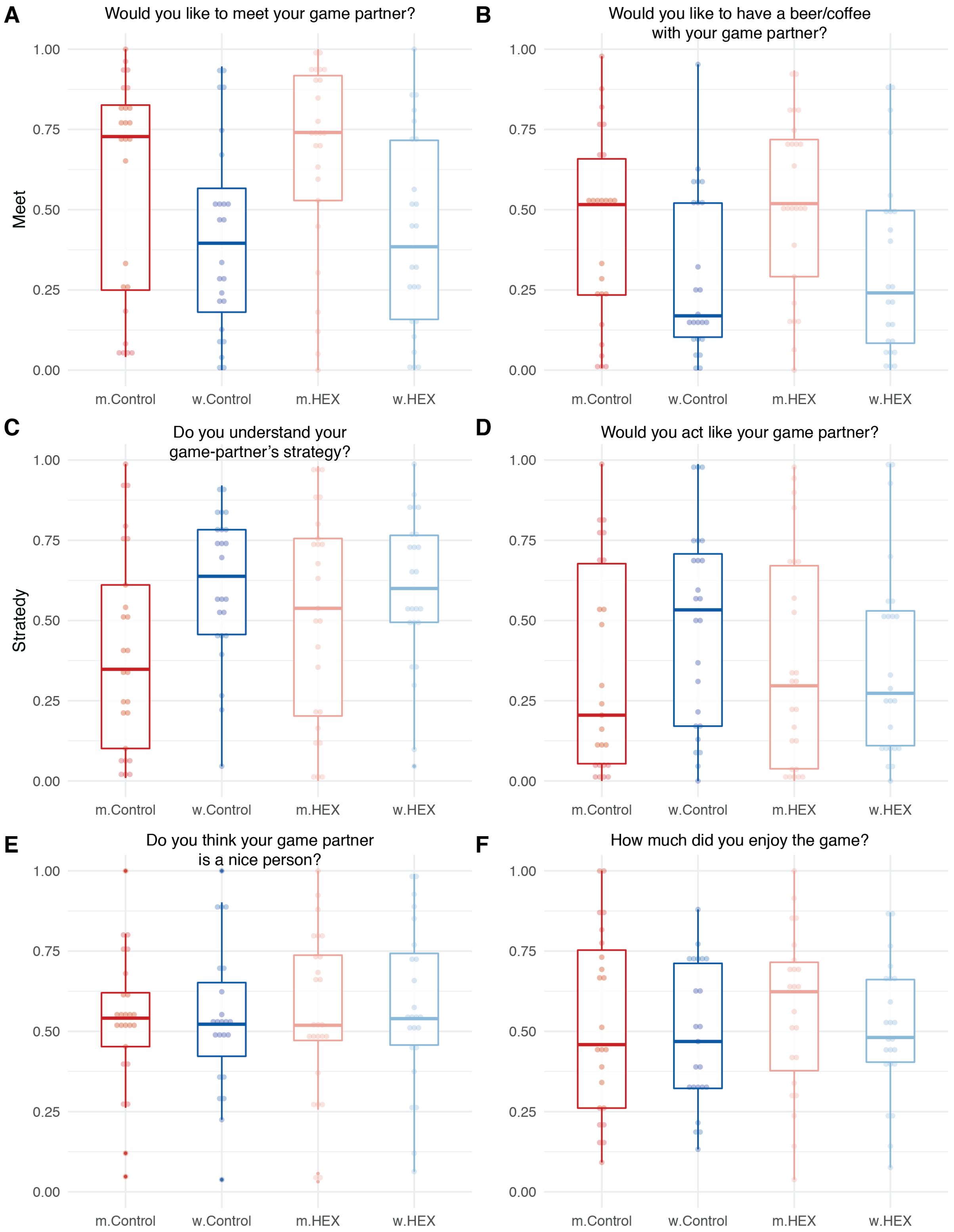

**Supplementary Fig. 9. Participants' attitudes towards their game-partner during the FC-PSAP**

(**A**) *Would you like to meet your game-partner?* Linear mixed model with factors of Odor, Sex and random effects of participant revealed a main effect of Sex (F(1) = 5.84, p = 0.02), no effect of Odor (F(1) = 1.17, p = 0.28) and no effect of the interaction Sex x Odor (F(1) = 0.77.

(**B**) *Would you like to have a beer or coffee with your game-partner?* Linear mixed model with factors of Odor, Sex and random effects of participant revealed main effect of Sex: $F(1) = 5.89$, $p = 0.02$, no effect of Odor: $F(1) = 0.03$, $p = 0.86$, and no effect of the interaction Sex x Odor: $F(1) = 0.12$, $p = 0.73$). (**C**) *Do you understand your game-partner's strategy?* Linear mixed model with factors of Odor, Sex and random effects of participant revealed a marginal effect of Sex: $F(1) = 3.82$, $p = 0.06$, no effect of Odor: $F(1) = 1.88$, $p = 0.18$, and a trend towards interaction of Sex x Odor: $F(1) = 3.14$, $p = 0.08$). Post-hoc analysis of the pairwise comparison revealed a trend towards a difference between women and men in the Control condition ($t(47) = 2.54$, $p = 0.06$). (**D**) *Would you act like your game partner did?* Linear mixed model with factors of Odor, Sex and random effects of participant revealed no effects (Sex: $F(1) = 0.80$, $p = 0.38$, Odor: $F(1) = 1.16$, $p = 0.29$, Sex x Odor: $F(1) = 2.58$, $p = 0.11$). (**E**) *Do you think your game-partner is a nice person?* Linear mixed model with factors of Odor, Sex and random effects of participant revealed no effects (Sex: $F(1) = 0.06$, $p = 0.80$, Odor: $F(1) = 0.49$, $p = 0.48$, Sex x Odor: $F(1) = 0.009$, $p = 0.92$). (**F**) *How much did you enjoy the game?* Linear mixed model with factors of Odor, Sex and random effects of participant revealed no effects (Sex ($F(1) = 0.72$, $p = 0.40$), no effect of Odor ($F(1) = 0.39$, $p = 0.54$, Sex x Odor ($F(1) = 0.06$, $p = 0.80$)).
